# Task-specific topology of brain networks supporting working memory and inhibition

**DOI:** 10.1101/2024.04.05.588287

**Authors:** Timofey Adamovich, Victoria Ismatullina, Nadezhda Chipeeva, Ilya Zakharov, Inna Feklicheva, Sergey Malykh

## Abstract

Network neuroscience investigates the brain’s connectome, revealing that cognitive functions are underpinned by dynamic neural networks. This study investigates how distinct cognitive abilities—working memory and inhibition—are supported by unique brain network configurations, which are constructed by estimating whole-brain networks through mutual information. The study involved 195 participants who completed the Sternberg Item Recognition and Flanker tasks while undergoing EEG recording. A mixed-effects linear model analyzed the influence of network metrics on cognitive performance, considering individual differences and task-specific dynamics. Results indicate that working memory and inhibition are associated with different network attributes, with working memory relying on distributed networks and inhibition on more segregated ones. Our analysis suggests that both strong and weak connections contribute to cognitive processes, as weak connections could potentially lead to a more stable and support networks of memory and inhibition. The findings indirectly support the Network Neuroscience Theory of Intelligence, suggesting different functional topology of networks inherent to various cognitive functions. Nevertheless, we propose that understanding individual variations in cognitive abilities requires recognizing both shared and unique processes within the brain’s network dynamics.

**Author summary:** This study analyzes how working memory and inhibition correspond to distinct neural network patterns by constructing whole-brain networks via mutual information from EEG data of 195 subjects performing cognitive tasks. Findings reveal working memory is supported by distributed connections while inhibition depends on segregated ones. The research underscores the importance of both strong and weak neural connections in cognitive function and supports the notion that cognitive functions emerge from the brain network of distinct topology. Moreover, it highlights the need to account for individual and task-specific variations to fully grasp the diverse network dynamics influencing cognitive abilities.

## Introduction

Network neuroscience combines graph theory and neuroimaging to decipher the brain’s intricate web of connections, illuminating how neural networks underpin cognitive functions by viewing the brain as a system of interlinked regions (Deco & Kringelbach, 2017; Sporns & Betzel, 2016). Research reveals that the brain operates as a small-world network, characterized by high clustering and short path lengths, facilitating both specialized (segregated) and holistic (integrated) processing (Cohen & D’Esposito, 2016; Deco et al., 2015). The brain’s dynamic network adapts to the cognitive demands of tasks like memory retrieval or problem-solving by reconfiguring connections to optimize function (Braun et al., 2015; Gonzalez-Castillo et al., 2015). Individual differences in network structure and adaptability are thought to contribute to cognitive performance variations, underscoring the importance of the brain’s dynamic reconfiguration in response to cognitive challenges (Cole et al., 2014; Finn et al., 2015; Krienen et al., 2014).

Investigation of the brain mechanisms of cognition must take into account a heterogeneous nature of cognitive functions. Theoretical models of intelligence, such as the Cattell-Horn-Carroll (CHC) theory, have been instrumental in conceptualizing intelligence as a hierarchical structure of abilities, ranging from general intelligence to broad and narrow cognitive abilities (Carroll, 1993; McGrew, 2009). However, the emergence of network neuroscience has provided a new lens through which to view cognition, linking these hierarchical models to the brain’s network architecture. The network neuroscience theory of intelligence (Barbey, 2018) posits that intelligence stems from the dynamic interplay of the brain’s networks, with the adaptability and flexibility of these networks being crucial for high-level cognitive functioning. According to this theory, different levels of intelligence correspond to the brain’s ability to reconfigure its networks. The theory further suggests that the manifestation of crystallized and fluid intelligence can be traced to the nature of connections within these networks—strong, stable connections and easy-to-reach brain states facilitate verbal intelligence, while weaker, more variable connections and hard-to-reach states might underpin nonverbal intelligence. The theory aligns with findings that associate efficient network reconfiguration with fluid intelligence, suggesting that the brain’s ability to transition between different network states in response to cognitive demands is a substrate of cognitive ability (Anderson & Barbey, 2023; Bassett et al., 2011; Dubois et al., 2018; Wen et al., 2015). Some recent studies, however, have found the opposite associations between brain reconfiguration and intelligence, namely the increased stability of brain networks was related to higher intelligence (Hilger et al., 2017, 2020; Pamplona et al., 2015; Thiele et al., 2022). Other studies were not able to establish a connection between general factor of intelligence on fMRI data of the Human Connectome Project (Kruschwitz et al., 2018) and attempted to replicate association between general intelligence and network measures on four independent datasets (Metzen et al., 2024).

In this study, we sought to expand on the premise that various types of intelligence are supported by distinct network underpinnings, by examining two cognitive abilities closely related to intelligence: working memory and inhibition.

Working memory (WM) (Furley & Memmert, 2010) and inhibition (Miyake & Friedman, 2012), two key cognitive abilities, are strongly linked to general intelligence (g). WM, crucial for reasoning and decision-making, has been shown to share about 50% of its variance with fluid intelligence (Ackerman et al., 2005; Kane et al., 2005). Conversely, the relationship between intelligence and inhibition, the capability to suppress distractions, is more contentious, with studies yielding mixed results across various ages (Duan & Shi, 2011; Friedman et al., 2006). Engle and Kane’s two-factor theory of cognitive control divides executive attention into goal maintenance and conflict resolution, highlighting their interaction in working memory capacity (Engle & Kane, 2004). Braver and colleagues (Braver et al., 2008) further dissect control processes in WM, including information selection, storage, updating, protection from interference, and usage to guide other cognitive systems. Conflict monitoring theory suggests that cognitive control is activated during tasks requiring response conflict resolution, with increased conflict leading to slower and less accurate responses.

Evidence for association between working memory, inhibition and functional connectivity are more robust than associations with general intelligence. Working memory primarily engages areas such as the prefrontal cortex, temporal, and parietal lobes, and patterns of functional connectivity could predict working memory performance (Avery et al., 2020; Finc et al., 2020; Murphy et al., 2020; Sala-Llonch et al., 2012). Furthermore, the neural underpinnings of working memory appear to be influenced by the nature of the task at hand; for instance, verbal working memory tasks typically activate the Broca’s area, while spatial working memory tasks draw on a more extensive network of brain regions (Bernal et al., 2015; Ren et al., 2019; Rogalsky et al., 2008).

Conversely, inhibition has been observed to involve neural networks and cognitive functions at multiple levels (Herstel & Wierenga, 2021). Cognitive control is typically linked to the activity within the cognitive control network (CCN), which includes the functioning of the prefrontal cortex, the dorsal anterior cingulate cortex, and the lateral prefrontal cortex. (Cole & Schneider, 2007). Several studies suggested that the reconfiguration of CCN is crucial for the effective filtering of information (Bartholomew et al., 2019; Cocchi et al., 2011; Marek et al., 2015; Menon & D’Esposito, 2022; Spielberg et al., 2015).

Drawing inspiration from the network neuroscience theory of intelligence (Barbey, 2018), we hypothesize that distinct cognitive processes may rely on different brain network configurations as well. Our study hypothesizes that the cognitive tasks of acquiring, maintaining, and updating information in working memory are underpinned by specific topological configurations of brain networks that differ from those supporting the inhibition of distracting stimuli.

## 2. Materials and methods

### 2.1. Sample

The participants were recruited through announcements in universities to participate in the study. There was no reward or payment. 195 participants took part in the study. Age ranged from 17 to 39 years (M = 20.3, SD = 3.35, 37% identified as female). Participants did not have any recorded history of psychiatric or neurological disorders and head traumas. All participants gave their consent to participate in the study. All procedure was approved by the Ethics Committee of the Psychological Institute of the Russian Academy of Education (protocol code 2019/2-10, date of approval 11.02.2019).

### 2.2. Cognitive tests

*Sternberg Item Recognition Paradigm (SIRP) or Sternberg Task* is a delayed stimulus identification test as a method for assessing working memory, which allows assessing working memory performance when solving problems of varying complexity. The test also allows distinguishing between such indicators as “search speed when performing a task” (mental scanning speed) and the direct success of completing a working memory task.

The test consists of 13 training and 72 test trials. In each trail, subjects are presented with a random set of numbers that they need to remember. The dial length varies from one to six digits, each of which is presented separately for 1.2 seconds. This is followed by a pause of 2 seconds, followed by a check digit. Subjects must press the right arrow button on the keyboard to answer “Yes, this is one of the memorized digits” or the left arrow button “No, this is a new digit” (correct and incorrect control digits are equally likely to appear), after which the control stimulus disappears, and the answer on the screen “Correct” or “Error” gives feedback on the correctness of the answer. The test has the following parameters: number of correct and incorrect answers, reaction time for each trial. For analysis Sternberg performance as number of correct answers (accuracy) and Sternberg RT as reaction time were used.

*Eriksen Flanker Task* is a set of response inhibition tasks and measures cognitive inhibition that measures the ability to inhibit responses when they are inappropriate for a given situational context. The participant presented with targeted stimuli along with distraction stimuli. Namely, an arrow pointing right or left, surrounded by other arrows. Congruent condition in the test — the target arrow is surrounded by other arrows pointing in the same direction. Incongruent condition — additional arrows point in the opposite direction. The participant’s task is to indicate in which direction the target arrow is pointing. A total of 200 test trials were presented, with half being congruent and the other half being incongruent. As part of the assessment of cognitive inhibition, only tasks for incongruent condition were used: number of correct answers (Flanker incongruent) and reaction time (Flanker incongruent RT). 200 test trials were presented, half congruent and half incongruent.

### 2.3. Procedure

The study involved two cognitive assessments: the Sternberg Item Recognition Test and the Flanker Task, which were administered in an alternating sequence. Each recording session started with a resting state. During resting state, participants were asked to relax, not to focus on anything specific, and to stay awake for 10 minutes. They received spoken commands every 2 minutes to either open or close their eyes. We only analyzed the EEG data collected while their eyes were open. Each cognitive task took from 15 to 20 minutes, so total time of the EEG acquisition was about 45-50 minutes.

The EEG data was recorded using a 64-electrode cap, set up according to the 10–10 international system, with a Brain Products ActiChamp amplifier. EEG was recorded in a sound-attenuated and electrically shielded room, with dim lighting. High conductive chloride gel was used to keep the impedance of the electrodes below 25 kOhm.

The EEG data were captured in real-time with the Brain Products PyCorder system, which recorded continuously at a 500 Hz sample rate without initially applying any filters. The reference point for the recording was the Cz electrode. During EEG-data preprocessing, we changed the reference to a REST reference (Yao, 2001) and reduced the sample rate to 256 Hz. Next, a high-pass filter at 1 Hz was applied. To clean the data, we used ICA decomposition to identify and remove typical interference like eye blinks and muscle movements. IClabel (Pion-Tonachini et al., 2019)was used to classify the components into brain and non-brain activity. Components marked as non-brain activity were removed from the data.

The EEG data was divided into 1-second chunks consequent for the resting state and lined up with stimuli presentation for task conditions. We used autoreject (Jas et al., 2017) to reject any segments containing artifacts. The data were considered fit for the study as long as less than 20% of epochs were flagged as bad across all the conditions.

### 2.4. EEG Analysis

Information transfer in the brain involves different frequency bands, each with unique characteristics. Accordingly, our investigation was performed in four distinct frequency ranges: theta (4-8 Hz), alpha (8-13 Hz), low beta (13-20 Hz), and high beta (20-30 Hz). The exploration of inter-frequency interactions was beyond the scope of this study. We present findings for each experimental condition across these specified frequency bands.

Previous studies (Bansal et al., 2019; J. Liu et al., 2018) observed a substantial inter-individual variability in functional connectivity. Studies also often focus on group-level findings, but there’s a growing recognition of the importance of individual differences in brain network measures. Studies have found that the degree of similarity between an individual’s brain connectivity in resting and active states can correlate with behavioral patterns, suggesting a personalized basis for intelligence (Tompson et al., 2018). This individual-specific manifestation of intelligence may be influenced by a person’s unique cognitive styles and traits.

In this study we wanted to accommodate for a possible spectrum of individual brain states associated with cognitive performance acknowledging the dynamic nature of brain activity. In order to achieve that we divide each condition into five non-overlapping intervals, each comprising 20% of relevant stimuli in the task starting with the first stimuli. The number of intervals was defined in order to consistently keep at least 10 stimuli in the interval. Within each interval the Gaussian-Copula mutual-information estimator (Ince et al., 2017) was used for calculation of functional connectivity for each stimulus that were then averaged within each interval to get the averaged connectivity matrix for the interval. Total of 5 connectivity matrices were constructed for each condition.

Adjacency matrices were transformed into networks through quartile thresholding (Adamovich et al., 2022; van den Heuvel et al., 2017). We used two different threshold values: 0.5, as previously applied in our research (Zakharov et al., 2020), and 0.8, a more commonly used threshold in the literature. Entries in each connectivity matrix falling below the respective quantile threshold were reassigned a value of zero. To preserve network integrity and avoid isolated nodes, any nodes that became disconnected post-thresholding were reconnected to their two strongest links.

Network structure was characterized using several metrics, including:

1. Average path length. The average path length is a measure of the average number of steps (or cost of transfer between nodes) along the shortest paths for all possible pairs of network nodes.
2. Betweenness centrality. Betweenness centrality quantifies the number of times a node acts as a bridge along the shortest path between two other nodes. Nodes with high betweenness centrality play a critical role in the flow of information within the network. Eigenvector centrality. Eigenvector centrality measures a node’s influence by considering the number and quality of its connections; nodes with high eigenvector centrality are connected to many nodes who themselves have high centrality.
3. Clustering coefficient. The clustering coefficient is a measure of the degree to which nodes in a network tend to cluster together. A higher clustering coefficient indicates a greater ‘local’ connectivity, which in EEG studies can suggest local processing or functional segregation within the brain. It reflects the presence of tightly-knit groups of regions that may specialize in specific cognitive functions.
4. Modularity. Modularity is a measure of the strength of division of a network into modules. Networks with high modularity have dense connections between the nodes within modules but sparse connections between nodes in different modules.
5. Participation coefficient. The participation coefficient quantifies how well a node interacts with multiple modules in the network. A high participation coefficient proposes that a node has many connections to nodes in different modules.
6. Rich club coefficient. The rich club coefficient measures the tendency of high-degree nodes (hubs) to be more interconnected than expected by chance. If a network has a high rich club coefficient, it suggests the presence of a ‘rich club’ – a cohesive group of central nodes that are highly interlinked and may coordinate complex network functions.

### 2.5. Statistical Analysis

Previous research has shown that information processing speed is an important predictor of cognitive success, along with the actual accuracy of the response (Meiran & Shahar, 2018; Schubert et al., 2017), so cognitive performance in the two tasks was quantified through accuracy and response time, yielding four behavioral metrics. The approach for estimating cognitive measures mirrored the connectivity analysis; each task was divided into five segments, with average accuracy and response time calculated for each segment, resulting in five data points per cognitive measure that corresponded to the connectivity matrices. For the Flanker task, data were exclusively drawn from incongruent trials. To achieve a more Gaussian distribution of response times, we applied a Box-Cox transformation.

The research aimed to discern the influence of connectivity on cognitive performance within the Sternberg and Flanker tasks. We employed a mixed-effects linear model (Baayen et al., 2008) with random intercepts to estimate the impact of all network metrics on cognitive outcomes, treating participant ID as a random effect and conducting the analysis for each frequency band separately. Prior to fitting the model, all variables were standardized. The Sternberg task was segmented into blocks, distinguishing between the set presentation (encoding phase) and the target number presentation (retrieval phase).

To improve the robustness of our findings, we implemented a bootstrap resampling method. Across 2000 iterations, the model was fit to a bootstrapped dataset, assembled by randomly sampling participants with replacement, maintaining the original dataset’s size. The resultant distribution of prediction coefficients is used as result of the analysis. A regression effect was deemed significant if the two-tailed 10% confidence interval of the distribution did not encompass zero.

We use MNE-python (Gramfort et al., 2014) for EEG preprocessing, the Frites software package (Combrisson et al., 2022) for estimation of functional connectivity, networkx (Hagberg et al., 2008) for graph measures calculation and lme4 (Bates et al., 2015) for modelling.

## 3. Results

First, Spearman’s correlation was used to assess the relationships between behavioral metrics (see table № 2). The correlation of behavioral characteristics corresponds to those previously presented in the literature.

**Table No 1.** Descriptive statistics for cognitive tests.

|  | **Sternberg**<br>**accuracy** | **Sternberg RT** | **Flanker**<br>**accuracy** | **Flanker RT** |
| --- | --- | --- | --- | --- |
| **Mean** | 0.901 | 0.922 | 0.355 | 0.794 |
| **Median** | 0.929 | 0.850 | 0.350 | 0.902 |
| **Standard**<br>**deviation** | 0.124 | 0.384 | 0.266 | 0.230 |

**Table No 2.** Correlation between cognitive measures.

|  | **Sternberg**<br>**accuracy** | **Sternberg RT** | **Flanker**<br>**accuracy** | **Flanker RT** |
| --- | --- | --- | --- | --- |
| **Sternberg**<br>**accuracy** | — |  |  |  |
| **Sternberg RT** | -0.098* | — |  |  |
| **Flanker**<br>**accuracy** | 0.052 | -0.240*** | — |  |
| **Flanker RT** | 0.073 | -0.021* | 0.199*** | — |

Note. \* $p < .05$ , \*\* $p < .01$ , \*\*\* $p < .001$ . Response times were box-cox transformed

The accuracy and response time for five segments of the test were calculated separately. Descriptive statistics for each of the five segments are presented in the table below (see Table 3).

**Table No 3.** Descriptive statistics for cognitive tasks per segments.

|  | Test segment | Sternberg accuracy | Sternberg RT | Flanker accuracy | Flanker RT |
| --- | --- | --- | --- | --- | --- |
| **Mean** | 1 | 0.882 | 1.03 | 0.278 | 0.725 |
|  | 2 | 0.909 | 0.886 | 0.374 | 0.795 |
|  | 3 | 0.902 | 0.902 | 0.373 | 0.815 |
|  | 4 | 0.903 | 0.922 | 0.363 | 0.812 |
|  | 5 | 0.907 | 0.873 | 0.387 | 0.826 |
| **Median** | 1 | 0.917 | 0.934 | 0.250 | 0.846 |
|  | 2 | 0.933 | 0.863 | 0.350 | 0.904 |
|  | 3 | 0.929 | 0.813 | 0.350 | 0.915 |
|  | 4 | 0.923 | 0.828 | 0.350 | 0.915 |
|  | 5 | 0.938 | 0.807 | 0.400 | 0.907 |
| **SD** | 1 | 0.140 | 0.428 | 0.220 | 0.253 |
|  | 2 | 0.126 | 0.255 | 0.284 | 0.235 |
|  | 3 | 0.114 | 0.418 | 0.276 | 0.219 |
|  | 4 | 0.117 | 0.479 | 0.268 | 0.226 |
|  | 5 | 0.122 | 0.270 | 0.265 | 0.203 |

To compare differences across test segments and ensure stability of the behavioral measures within the test, a One-Way Repeated Measures ANOVA was conducted (see Table 4).

**Table No 4.**
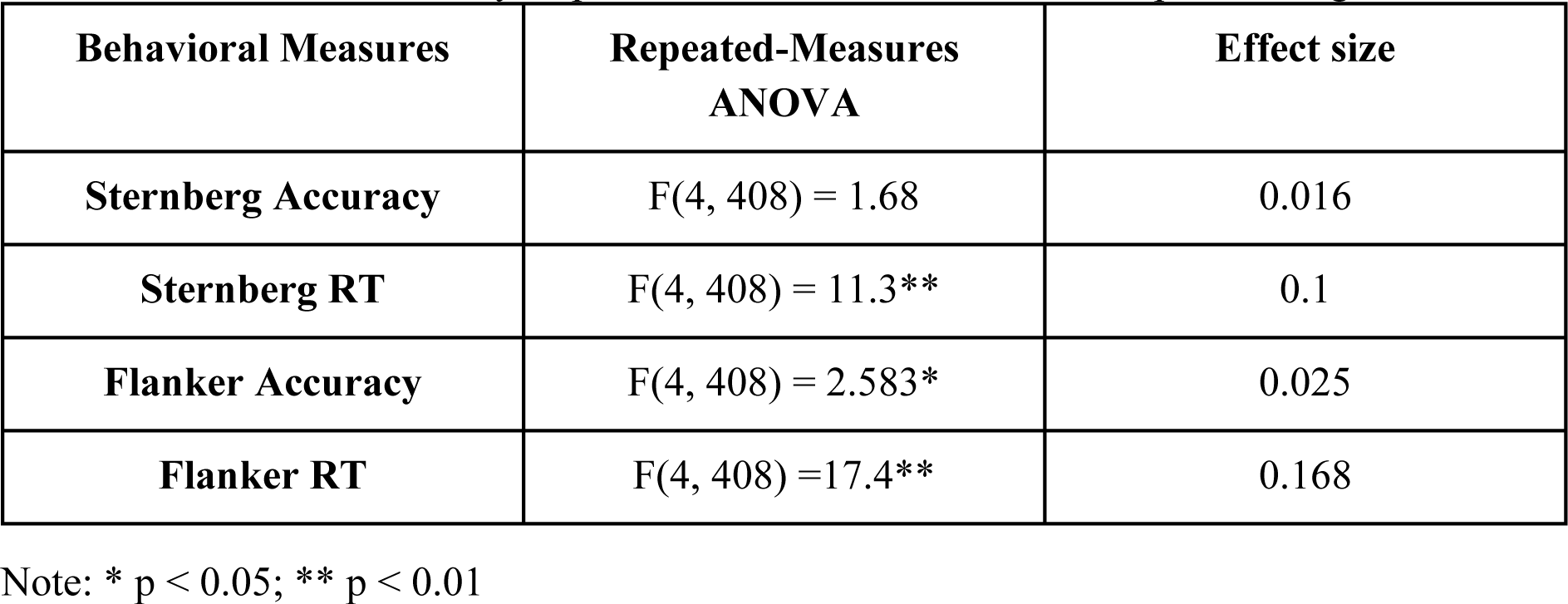
Results of One-Way Repeated Measures ANOVA for five parts of cognitive tasks.

| Behavioral Measures | Repeated-Measures ANOVA | Effect size |
| --- | --- | --- |
| **Sternberg Accuracy** | $F(4, 408) = 1.68$ | 0.016 |
| **Sternberg RT** | $F(4, 408) = 11.3^{**}$ | 0.1 |
| **Flanker Accuracy** | $F(4, 408) = 2.583^*$ | 0.025 |
| **Flanker RT** | $F(4, 408) = 17.4^{**}$ | 0.168 |

Note: \* $p < 0.05$ ; \*\* $p < 0.01$

One-Way ANOVA indicates the differences in the behavioral score related to poorer performance in the first 20% of the Flanker accuracy and response time in both tasks. During the rest of the test the performance was stable.

The objective of this article is to identify common and unique topological features of functional connectivity that correlate with cognitive performance success across various cognitive functions. To achieve this goal, we used functional connectivity measured during task execution to predict cognitive task success within that framework, revealing the topological characteristics of networks associated with that cognitive function. Additionally, we determined the topological features of resting-state networks and their relationship with cognitive task success, assessing the influence of the brain’s background information exchange processes on performance under cognitive load. We also explored differences between these states to determine how the reconfiguration between a resting state and a state of cognitive load affects cognitive performance success.

### The effect of task-induced functional connectivity on performance in cognitive tasks

The aim of this section is to describe the most robust network characteristics estimated during the task associated with the success of executing a task. The results reported in this section are presented in Table 5 and Figure 1.

**Fig 1.**
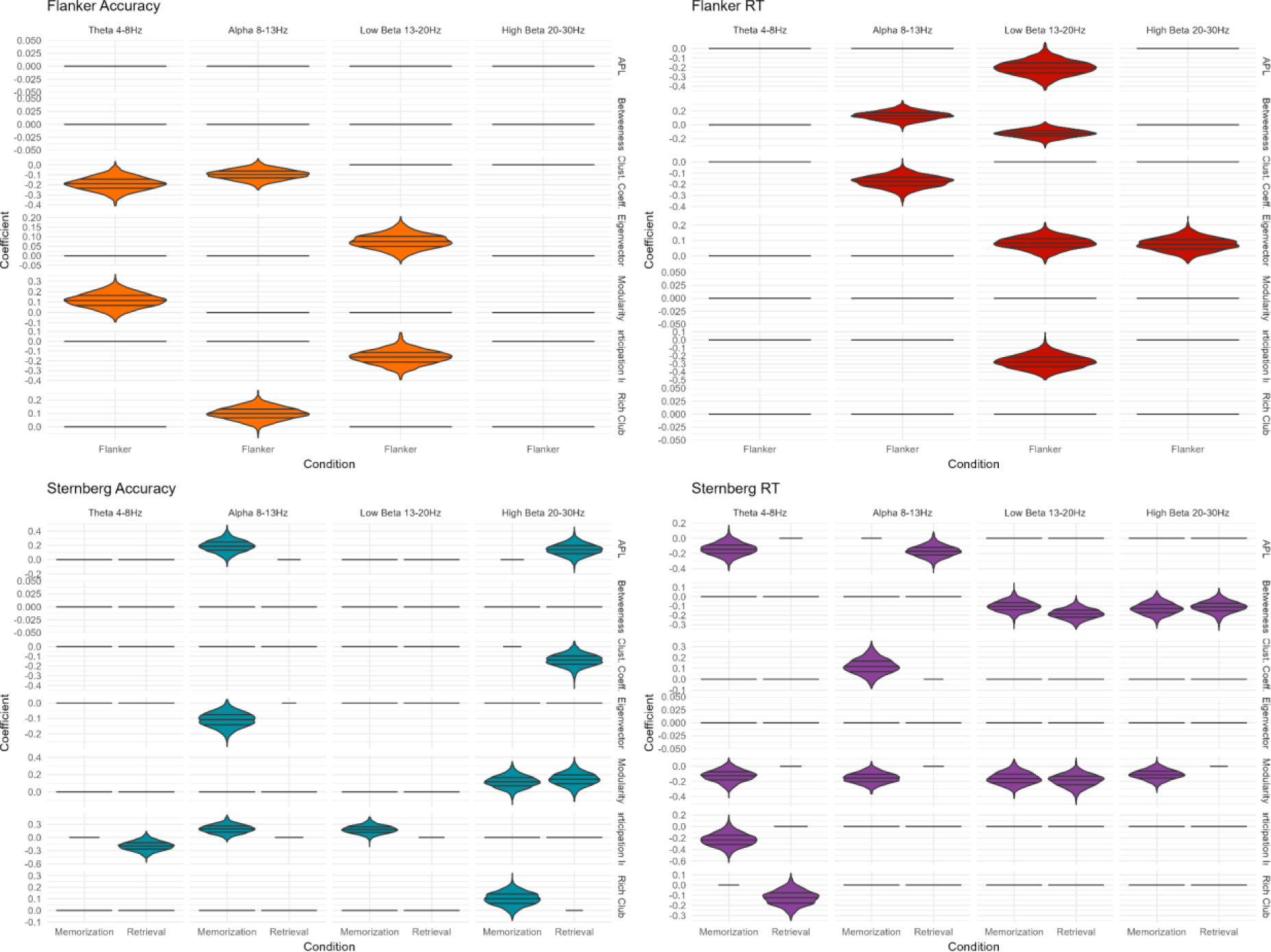
The significant effects of network measures during cognitive tasks on behavioral measures. The x-axis represents the EEG condition, while the y-axis displays the regression coefficients of network measures on behavioral performance. The frequencies are visualized column wise, and the network measures are arranged rowwise. Each plot corresponds to a specific behavioral measure, with the violin plot indicating the distribution’s 0.25, 0.5, and 0.75 quantiles. Horizontal lines indicates absence of significant effect and are set to 0.

**Table No 5.**
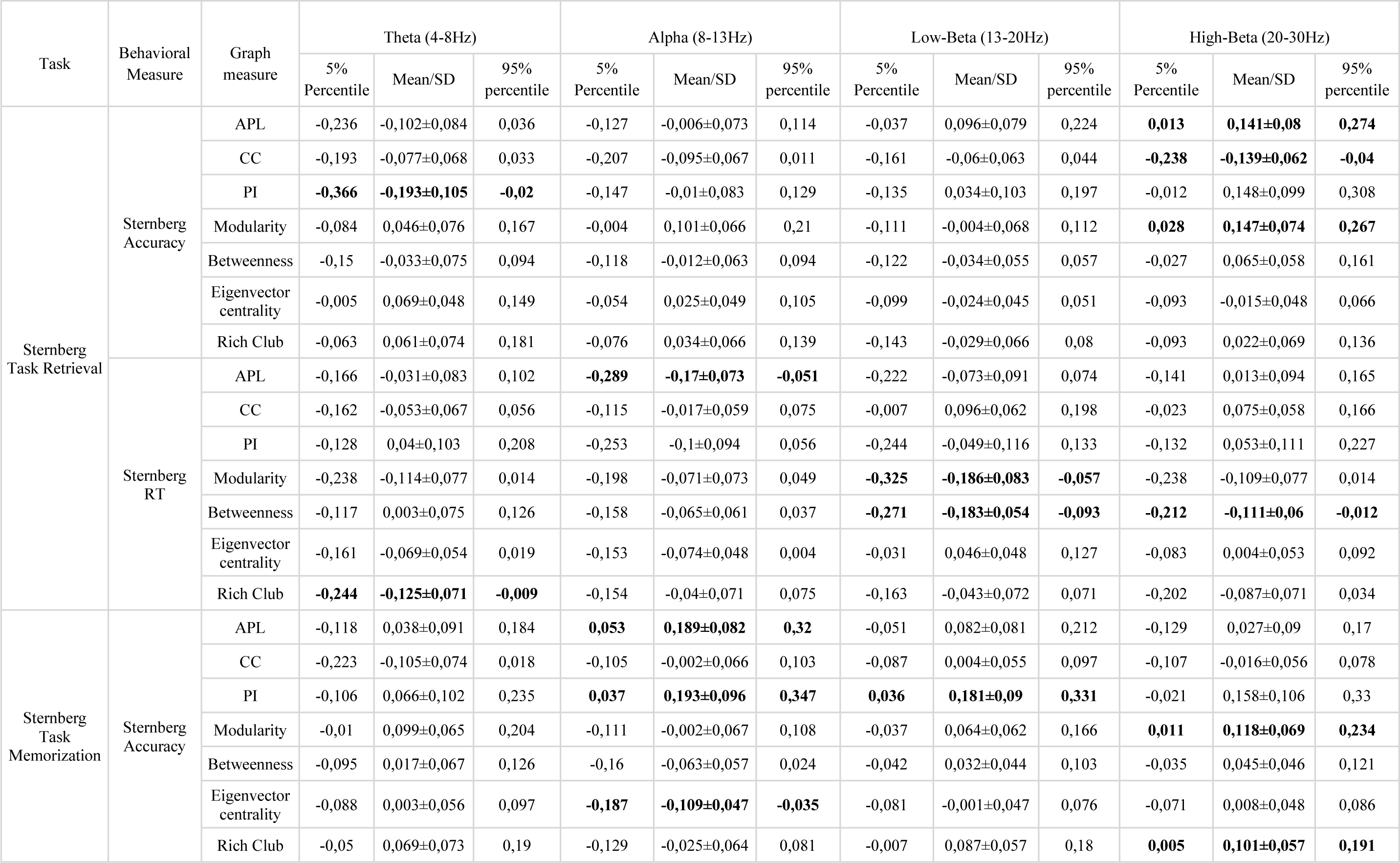
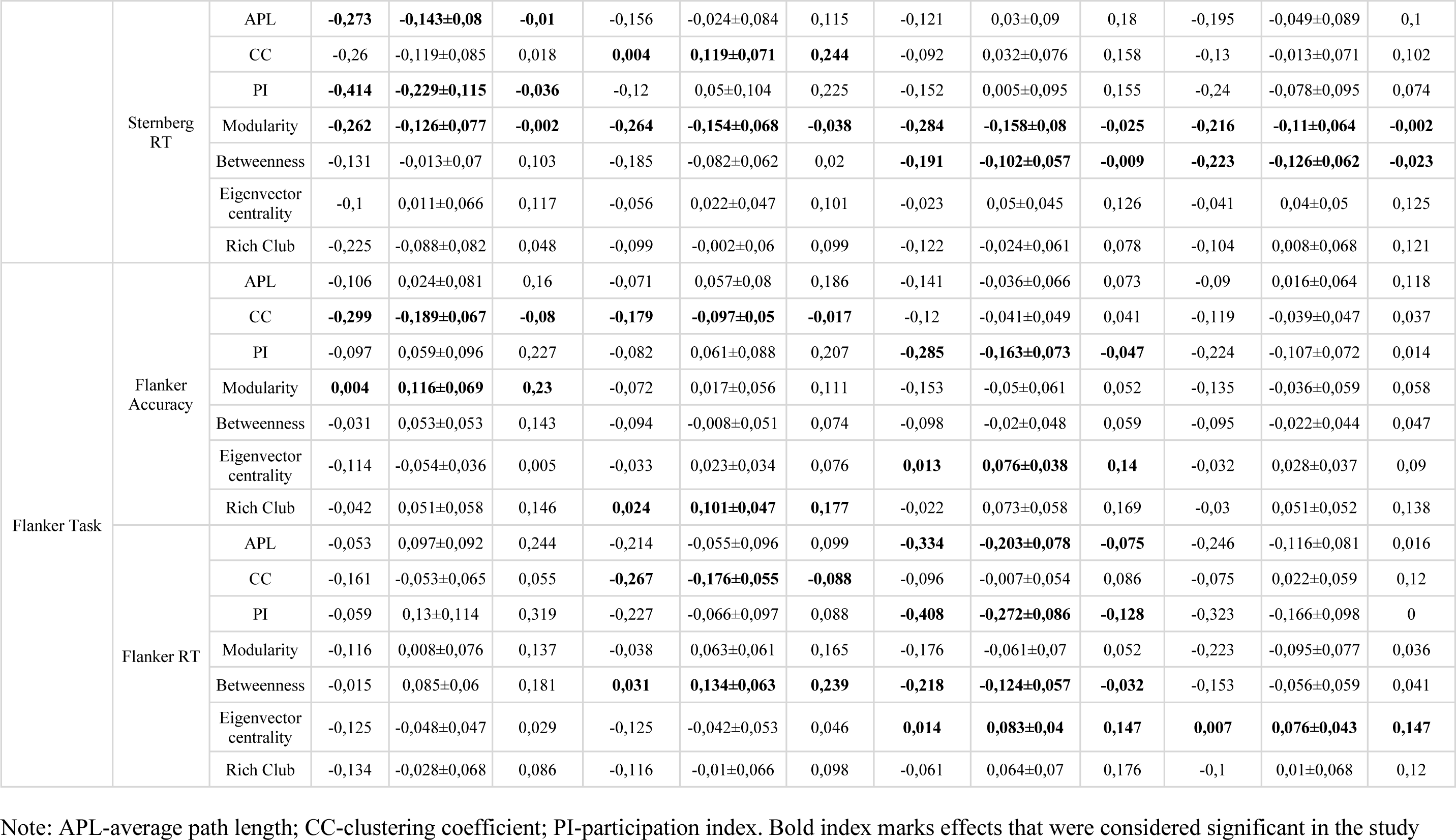
Regression coefficients for cognitive tasks.

The relationship between path length and execution time (inversely) in alpha and theta bands is the most significant. In the alpha band, path length during the memorization phase is positively associated with task execution accuracy. A similar relationship is observed in the high beta range, suggesting that increased information exchange between different brain areas leads to decreased response time and improved task performance. Betweenness centrality is predominantly associated with response time in low and high beta bands, both during the memorization and retrieval phases. Reduced centrality levels are linked to increased response times. The clustering coefficient does not show a clear connection with the Sternberg task, except for a positive relationship during the memorization phase in high beta range. Response time is positively related to the clustering coefficient in the alpha range. Eigenvector centrality during memorization has a negative relation to response time in the alpha band. Modularity has clear negative relationships with response time during the memorization phase across all bands and additionally with retrieval phase in the low beta band. Task accuracy is associated with modularity in the high beta band for both phases. Increased modularity is associated with faster and more successful task performance. The participation index has opposite associations. In the alpha and low beta bands an increase in the index is related to more successful task performance, while a decrease in theta band is associated with better performance. The rich club coefficient - a lower value of this indicator in theta band leads to longer execution time, while an increase in the memorization phase is associated with a higher level of success.

No significant associations of the Flanker task with path length were found. Betweenness centrality is inversely associated with response time, as higher betweenness in alpha band and lower in the lower beta band are related to higher response time. Participants with higher clustering coefficient values in the alpha range and theta range have lower accuracy and slower response time. An increase in eigenvector centrality value in the low beta range is associated with higher performance success and execution time. Similar associations are also observed in high beta bands but only with execution time. An increase in modularity in the theta rhythm is associated with more successful task execution. A decrease in the participation index in the alpha range is associated with reduced success and increased reaction time. The rich-club index is associated with task execution accuracy only in the alpha range.

We can draw several conclusions regarding the cognitive processes involved in Sternberg and Flanker tasks. In the Sternberg task, accuracy is related to brain network dynamics within frequency bands, especially in the alpha and beta bands. The path length, modularity, and participation index within these bands emerge as significant factors, reflecting the importance of efficient global network integration and the balance between distributed and specialized processing for accurate task performance. Response time in the Sternberg task, however, appears less tied to specific frequency bands, indicating a broader reliance on the brain’s overall network properties. Here, betweenness centrality and modularity are highlighted, indicating the need for a well-organized network structure that can quickly adapt and reconfigure to meet the task demands. Notably, these network characteristics during the memorization phase seem to be a better predictor of both accuracy and response times.

In the Flanker task the patterns of brain network associations with cognitive performance differ markedly from the Sternberg task. For accuracy, lower frequencies are implicated, with the clustering coefficient, modularity, and rich club coefficient being influential. This suggests that at lower frequencies, a balance between local processing and integration is essential for correct inhibition of irrelevant information. The response time in the Flanker task, conversely, is associated with higher frequency ranges and hinges primarily on measures of centrality. This points to the speed of processing and the agility of the brain’s information routing being facilitated by central nodes that likely play a pivotal role in rapid attention shifting and response execution.

Differences in the associations between network measures with accuracy and response time, observed in both tasks, may suggest that the brain mechanisms associated with these behavioral characteristics are distinct, and the characteristics themselves are not equivalent but relatively independent, each relating to different aspects of cognition.

### The effect of resting-state functional connectivity on performance in cognitive tasks

The second section focuses on the evaluation of the brain’s intrinsic characteristics of information exchange as predictors of success in cognitive tasks, based on resting EEG data. The results reported in this section are presented in Table 7 and Figure 2. It has been found that the path length between brain regions does not significantly correlate with the performance in Sternberg and Flanker tasks. However, the betweenness centrality coefficient shows a unique relationship with the success rate of the Flanker task in alpha band. The clustering coefficient demonstrates a connection with performance in both tasks: for the Flanker task, it correlates with both the accuracy in both beta bands and the response time in alpha band and low beta band, while in the Sternberg task, it is linked with the performance time and accuracy in alpha band frequency. Eigenvector centrality is also a significant predictor of performance across tasks; in the Flanker task, it relates to the response time in theta band and alpha band, and to the accuracy in alpha band and both beta bands. In the Sternberg task, response time is associated with eigenvector centrality in alpha band and high beta. Modularity corresponds to accuracy in the Sternberg task in theta band and to both accuracy and response time in the Flanker task within the alpha band. Additionally, the participation index is exclusively tied to the accuracy of the Flanker task in low beta band. Lastly, the response time for the Sternberg task at various frequencies (theta, alpha and both beta bands) is linked with the brain’s rich club network, indicating that nodes with more extensive connections to other nodes in the network are associated with longer performance times. Thus, we have two distinct sets of network measures related to performance in two cognitive tasks.

**Fig 2.**
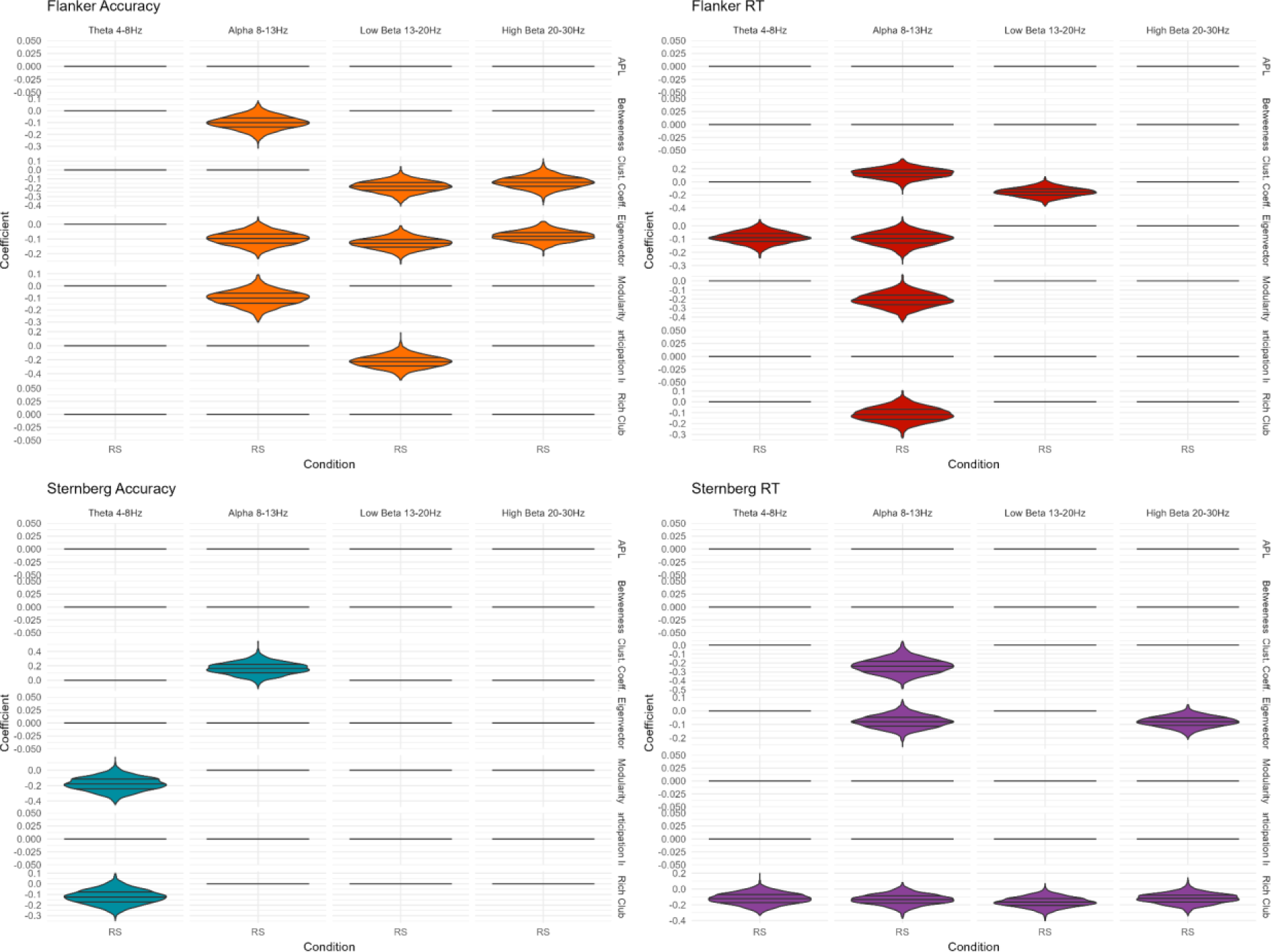
The significant effects of resting state network measures on behavioral measures. The x-axis represents the EEG condition, while the y-axis displays the regression coefficients of network measures on behavioral performance. The frequencies are visualized column wise, and the network measures are arranged rowwise. Each plot corresponds to a specific behavioral measure, with the violin plot indicating the distribution’s 0.25, 0.5, and 0.75 quantiles. Horizontal lines indicate absence of significant effect and were set to 0.

**Table # 7.**
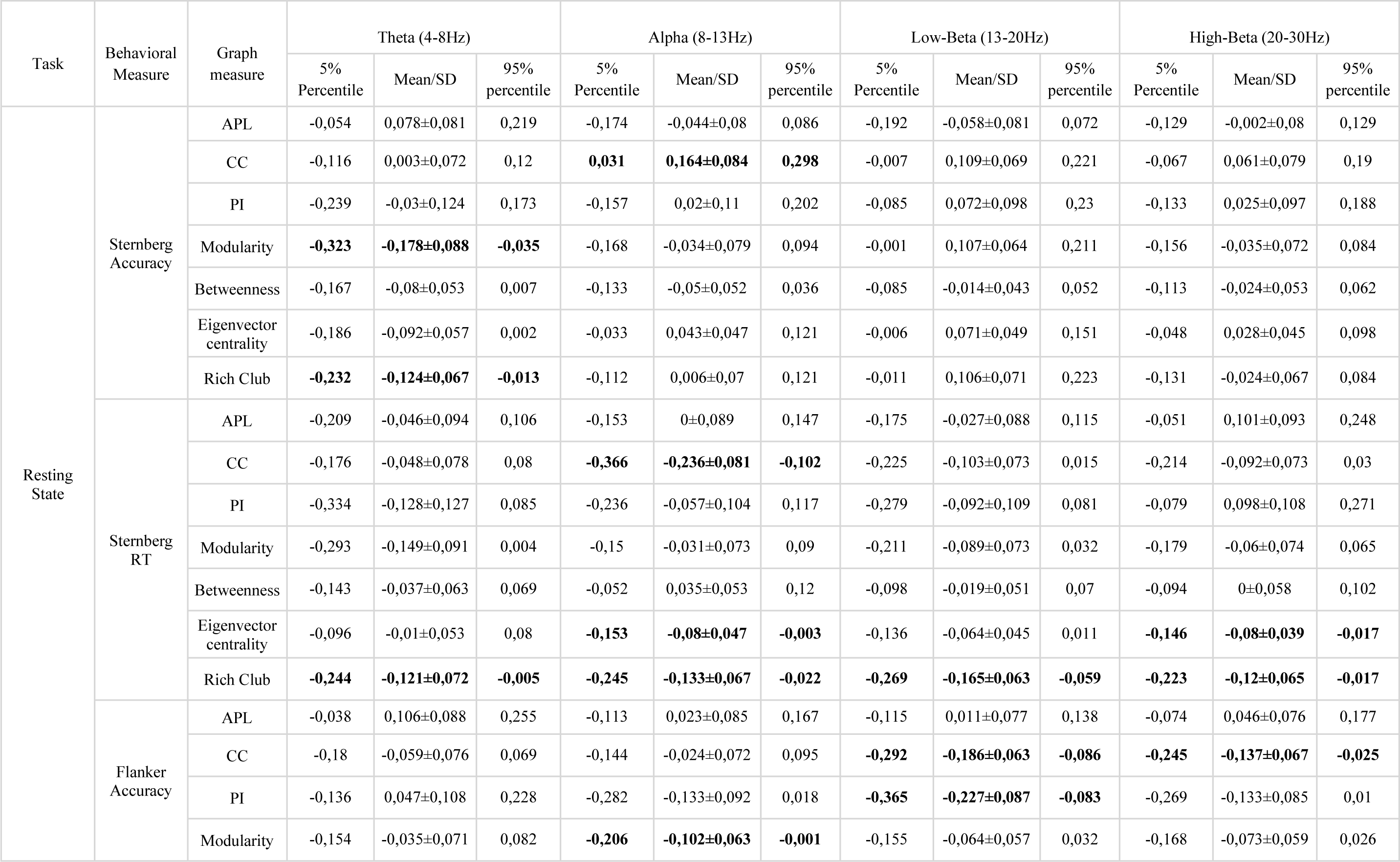
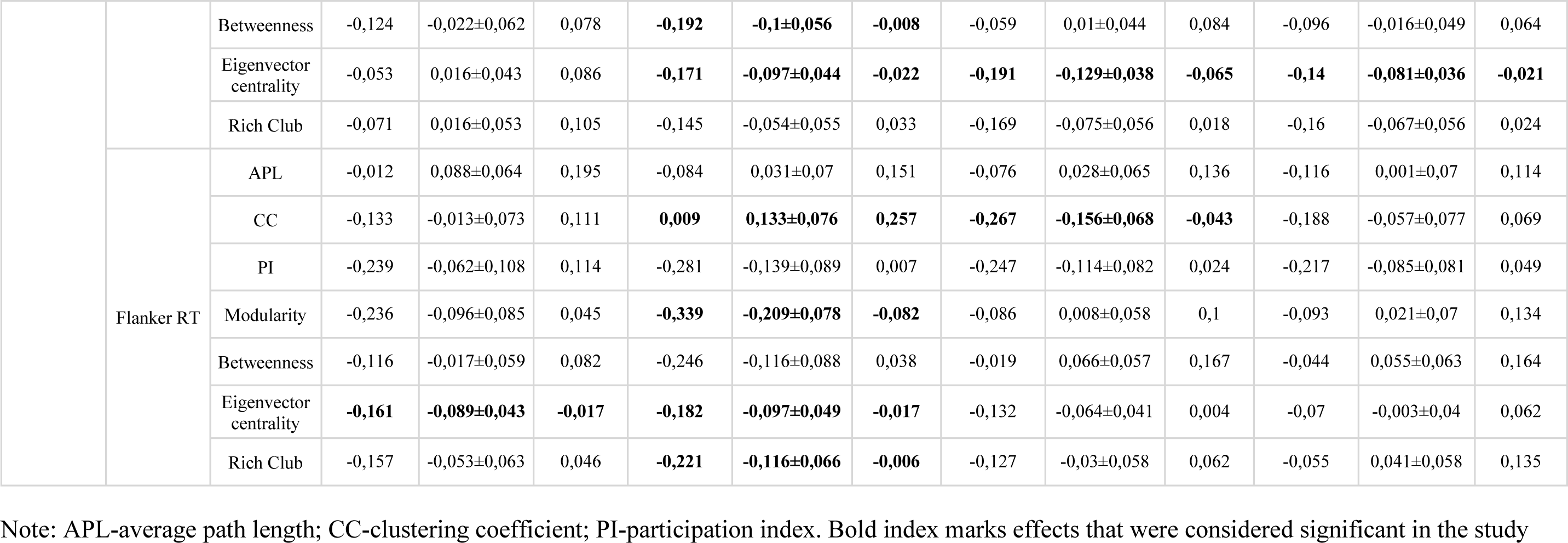
Regression coefficients for the resting state condition.

| Task | Behavioral Measure | Graph measure | Theta (4-8Hz) |  |  | Alpha (8-13Hz) |  |  | Low-Beta (13-20Hz) |  |  | High-Beta (20-30Hz) |  |  |
| --- | --- | --- | --- | --- | --- | --- | --- | --- | --- | --- | --- | --- | --- | --- |
|  |  |  | 5% Percentile | Mean/SD | 95% percentile | 5% Percentile | Mean/SD | 95% percentile | 5% Percentile | Mean/SD | 95% percentile | 5% Percentile | Mean/SD | 95% percentile |
| Resting State | Sternberg Accuracy | APL | -0,054 | 0,078±0,081 | 0,219 | -0,174 | -0,044±0,08 | 0,086 | -0,192 | -0,058±0,081 | 0,072 | -0,129 | -0,002±0,08 | 0,129 |
|  |  | CC | -0,116 | 0,003±0,072 | 0,12 | **0,031** | **0,164±0,084** | **0,298** | -0,007 | 0,109±0,069 | 0,221 | -0,067 | 0,061±0,079 | 0,19 |
|  |  | PI | -0,239 | -0,03±0,124 | 0,173 | -0,157 | 0,02±0,11 | 0,202 | -0,085 | 0,072±0,098 | 0,23 | -0,133 | 0,025±0,097 | 0,188 |
|  |  | Modularity | **-0,323** | **-0,178±0,088** | **-0,035** | -0,168 | -0,034±0,079 | 0,094 | -0,001 | 0,107±0,064 | 0,211 | -0,156 | -0,035±0,072 | 0,084 |
|  |  | Betweenness | -0,167 | -0,08±0,053 | 0,007 | -0,133 | -0,05±0,052 | 0,036 | -0,085 | -0,014±0,043 | 0,052 | -0,113 | -0,024±0,053 | 0,062 |
|  |  | Eigenvector centrality | -0,186 | -0,092±0,057 | 0,002 | -0,033 | 0,043±0,047 | 0,121 | -0,006 | 0,071±0,049 | 0,151 | -0,048 | 0,028±0,045 | 0,098 |
|  |  | Rich Club | **-0,232** | **-0,124±0,067** | **-0,013** | -0,112 | 0,006±0,07 | 0,121 | -0,011 | 0,106±0,071 | 0,223 | -0,131 | -0,024±0,067 | 0,084 |
|  | Sternberg RT | APL | -0,209 | -0,046±0,094 | 0,106 | -0,153 | 0±0,089 | 0,147 | -0,175 | -0,027±0,088 | 0,115 | -0,051 | 0,101±0,093 | 0,248 |
|  |  | CC | -0,176 | -0,048±0,078 | 0,08 | **-0,366** | **-0,236±0,081** | **-0,102** | -0,225 | -0,103±0,073 | 0,015 | -0,214 | -0,092±0,073 | 0,03 |
|  |  | PI | -0,334 | -0,128±0,127 | 0,085 | -0,236 | -0,057±0,104 | 0,117 | -0,279 | -0,092±0,109 | 0,081 | -0,079 | 0,098±0,108 | 0,271 |
|  |  | Modularity | -0,293 | -0,149±0,091 | 0,004 | -0,15 | -0,031±0,073 | 0,09 | -0,211 | -0,089±0,073 | 0,032 | -0,179 | -0,06±0,074 | 0,065 |
|  |  | Betweenness | -0,143 | -0,037±0,063 | 0,069 | -0,052 | 0,035±0,053 | 0,12 | -0,098 | -0,019±0,051 | 0,07 | -0,094 | 0±0,058 | 0,102 |
|  |  | Eigenvector centrality | -0,096 | -0,01±0,053 | 0,08 | **-0,153** | **-0,08±0,047** | **-0,003** | -0,136 | -0,064±0,045 | 0,011 | **-0,146** | **-0,08±0,039** | **-0,017** |
|  |  | Rich Club | **-0,244** | **-0,121±0,072** | **-0,005** | **-0,245** | **-0,133±0,067** | **-0,022** | **-0,269** | **-0,165±0,063** | **-0,059** | **-0,223** | **-0,12±0,065** | **-0,017** |
|  | Flanker Accuracy | APL | -0,038 | 0,106±0,088 | 0,255 | -0,113 | 0,023±0,085 | 0,167 | -0,115 | 0,011±0,077 | 0,138 | -0,074 | 0,046±0,076 | 0,177 |
|  |  | CC | -0,18 | -0,059±0,076 | 0,069 | -0,144 | -0,024±0,072 | 0,095 | **-0,292** | **-0,186±0,063** | **-0,086** | **-0,245** | **-0,137±0,067** | **-0,025** |
|  |  | PI | -0,136 | 0,047±0,108 | 0,228 | -0,282 | -0,133±0,092 | 0,018 | **-0,365** | **-0,227±0,087** | **-0,083** | -0,269 | -0,133±0,085 | 0,01 |
|  |  | Modularity | -0,154 | -0,035±0,071 | 0,082 | **-0,206** | **-0,102±0,063** | **-0,001** | -0,155 | -0,064±0,057 | 0,032 | -0,168 | -0,073±0,059 | 0,026 |
|  |  | Betweenness | -0,124 | -0,022±0,062 | 0,078 | **-0,192** | **-0,1±0,056** | **-0,008** | -0,059 | 0,01±0,044 | 0,084 | -0,096 | -0,016±0,049 | 0,064 |
|  |  | Eigenvector centrality | -0,053 | 0,016±0,043 | 0,086 | **-0,171** | **-0,097±0,044** | **-0,022** | **-0,191** | **-0,129±0,038** | **-0,065** | **-0,14** | **-0,081±0,036** | **-0,021** |
|  |  | Rich Club | -0,071 | 0,016±0,053 | 0,105 | -0,145 | -0,054±0,055 | 0,033 | -0,169 | -0,075±0,056 | 0,018 | -0,16 | -0,067±0,056 | 0,024 |
|  | Flanker RT | APL | -0,012 | 0,088±0,064 | 0,195 | -0,084 | 0,031±0,07 | 0,151 | -0,076 | 0,028±0,065 | 0,136 | -0,116 | 0,001±0,07 | 0,114 |
|  |  | CC | -0,133 | -0,013±0,073 | 0,111 | **0,009** | **0,133±0,076** | **0,257** | **-0,267** | **-0,156±0,068** | **-0,043** | -0,188 | -0,057±0,077 | 0,069 |
|  |  | PI | -0,239 | -0,062±0,108 | 0,114 | -0,281 | -0,139±0,089 | 0,007 | -0,247 | -0,114±0,082 | 0,024 | -0,217 | -0,085±0,081 | 0,049 |
|  |  | Modularity | -0,236 | -0,096±0,085 | 0,045 | **-0,339** | **-0,209±0,078** | **-0,082** | -0,086 | 0,008±0,058 | 0,1 | -0,093 | 0,021±0,07 | 0,134 |
|  |  | Betweenness | -0,116 | -0,017±0,059 | 0,082 | -0,246 | -0,116±0,088 | 0,038 | -0,019 | 0,066±0,057 | 0,167 | -0,044 | 0,055±0,063 | 0,164 |
|  |  | Eigenvector centrality | **-0,161** | **-0,089±0,043** | **-0,017** | **-0,182** | **-0,097±0,049** | **-0,017** | -0,132 | -0,064±0,041 | 0,004 | -0,07 | -0,003±0,04 | 0,062 |
|  |  | Rich Club | -0,157 | -0,053±0,063 | 0,046 | **-0,221** | **-0,116±0,066** | **-0,006** | -0,127 | -0,03±0,058 | 0,062 | -0,055 | 0,041±0,058 | 0,135 |

Note: APL-average path length; CC-clustering coefficient; PI-participation index. Bold index marks effects that were considered significant in the study

Unique processes for the Sternberg task include:

1. Clustering Coefficient: The accuracy of the Sternberg task performance is related to the clustering coefficient in alpha band. A higher clustering coefficient is associated with a higher likelihood of correctly performing the task.
2. Eigenvector centrality: in high beta band lower eigenvector centrality is related to lower response time.
3. Modularity: Modularity is associated with performance in theta band only. A decrease in modularity is linked with more successful task completion.
4. Rich club coefficient: rich club coefficient is inversely associated with measures of task performance. This association spans across all frequency bands for response time and with accuracy in theta band only.

Unique processes for the Flanker task include:

1. Betweenness centrality: lower levels of betweenness centrality in alpha band are associated with the higher accuracy of the Flanker task
2. Clustering Coefficient: The accuracy of the Flanker task performance is related to the clustering coefficient in both beta bands. A higher clustering coefficient is associated with a higher likelihood of correctly performing the task. Association with response

time was found in alpha band and low beta band and is opposite for these bands. Higher clustering in alpha and lower clustering in beta is associated with higher response times.

1. 3. Eigenvector centrality: The accuracy is inversely associated with eigenvector centrality in the alpha band and both beta bands. Same association exists for response time in theta and alpha range.
2. 4. Modularity: Lower level of modularity in networks is associated with higher accuracy and higher response time in the Flanker task.
3. 5. Participation Index: The participation index is only associated with the accuracy of the Flanker task performance in low beta band.
4. 6. Rich club coefficient: Alpha band rich club coefficient is inversely associated with Flanker task response time.

Internal processes that are not directly linked to cognitive load, but rather believed to be rooted in the overall architecture of the brain, exhibit a stronger correlation with the Flanker task compared to the Sternberg task. The accuracy of responses in the Flanker task is linked to network characteristics in the alpha and both beta frequency bands, whereas the accuracy of responses in the Sternberg task is predominantly associated with network measures in the theta and alpha frequency bands, which is the opposite of what was observed during the task itself.

While there are no overlapping network measures for accuracy between the two tasks, a commonality is observed in the processes determining response times: eigenvector centrality and clustering coefficient in the alpha band. However, the direction of their influence diverges; a decrease in eigenvector centrality is generally associated with longer response times in both tasks, whereas the clustering coefficient’s impact varies—a higher clustering coefficient is linked with slower responses in the Flanker task, but in the Sternberg task, a lower clustering coefficient relates to longer response times.

For the Flanker task, the associations with response time predominantly involve a variety of network measures within the alpha band, underscoring the band’s importance in task-related processing speed. On the other hand, in the Sternberg task, the most robust association with response time is observed with the level of rich-club organization and spans across frequency bands, indicating that the integration of high-degree nodes into a cohesive core is vital for rapid memory processing.

Comparing the networks of the resting state and during the cognitive tasks, each behavioral measure exposes a unique pattern of network activity associated with performance. The efficacy of inhibitory control, as assessed in tasks like the Flanker, may be predicated on the architecture of these pre-existing networks, suggesting that the brain’s baseline state sets the stage for its ability to filter and suppress distractions. Conversely, the performance in working memory tasks, such as the Sternberg, seems to hinge on the brain’s agility to rapidly reconfigure its networks, aligning them with the immediate requirements of the task at hand.

Some knowledge could also be gathered from the changes in frequency specificity of the resting state and cognitive task conditions. The cognitive functioning suggests an increase in higher frequency neural oscillations during task engagement, which is indicative of the brain’s active information processing. However, a shift is observed in the context of the Flanker test, where accuracy correlates more significantly with lower frequency bands.

### Effects of reconfiguration between resting-state and task-induced functional connectivity on performance in cognitive tasks

To calculate the reconfiguration of networks between the resting state and the cognitive load state, the resting state value was subtracted from the corresponding metric obtained during the cognitive load. An increase in values along the scale indicates that the metric value of the networks is higher within the task with cognitive load. The purpose of this section is to assess the reconfiguration between the two states as a factor of success in cognitive tasks. The results reported in this section are presented in Table 8 and Figure 3.

**Fig 3.**
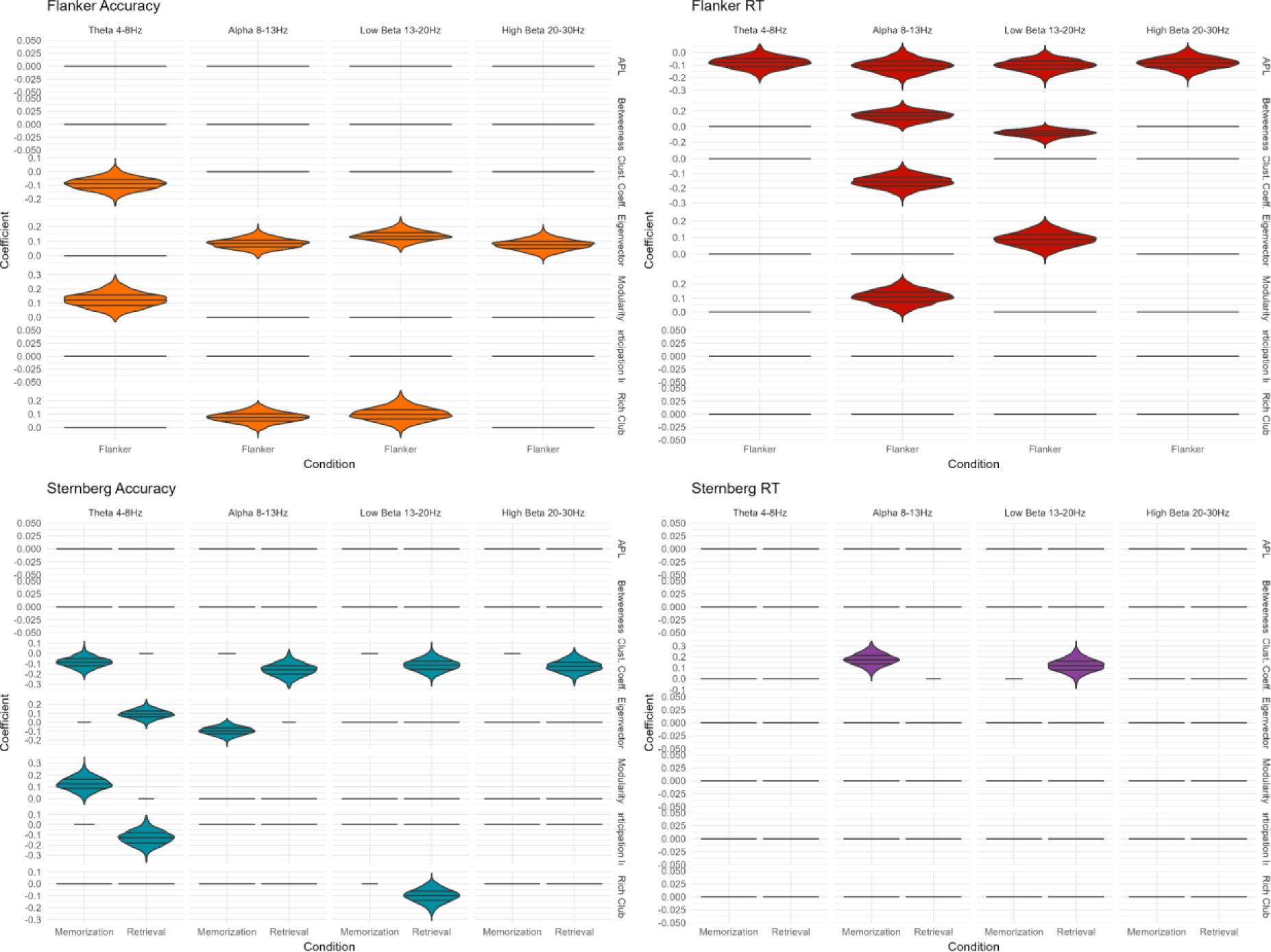
The significant effects of reconfiguration between resting state and cognitive state on behavioral measures. The x-axis represents the EEG condition, while the y-axis displays the regression coefficients of network measures on behavioral performance. The frequencies are visualized columnwise, and the network measures are arranged rowwise. Each plot corresponds to a specific behavioral measure, with the violin plot indicating the distribution’s 0.25, 0.5, and 0.75 quantiles. Horizontal lines indicate absence of significant effect and were set to 0. Horizontal lines indicate absence of significant effect and are set to 0.

**Fig 3.**
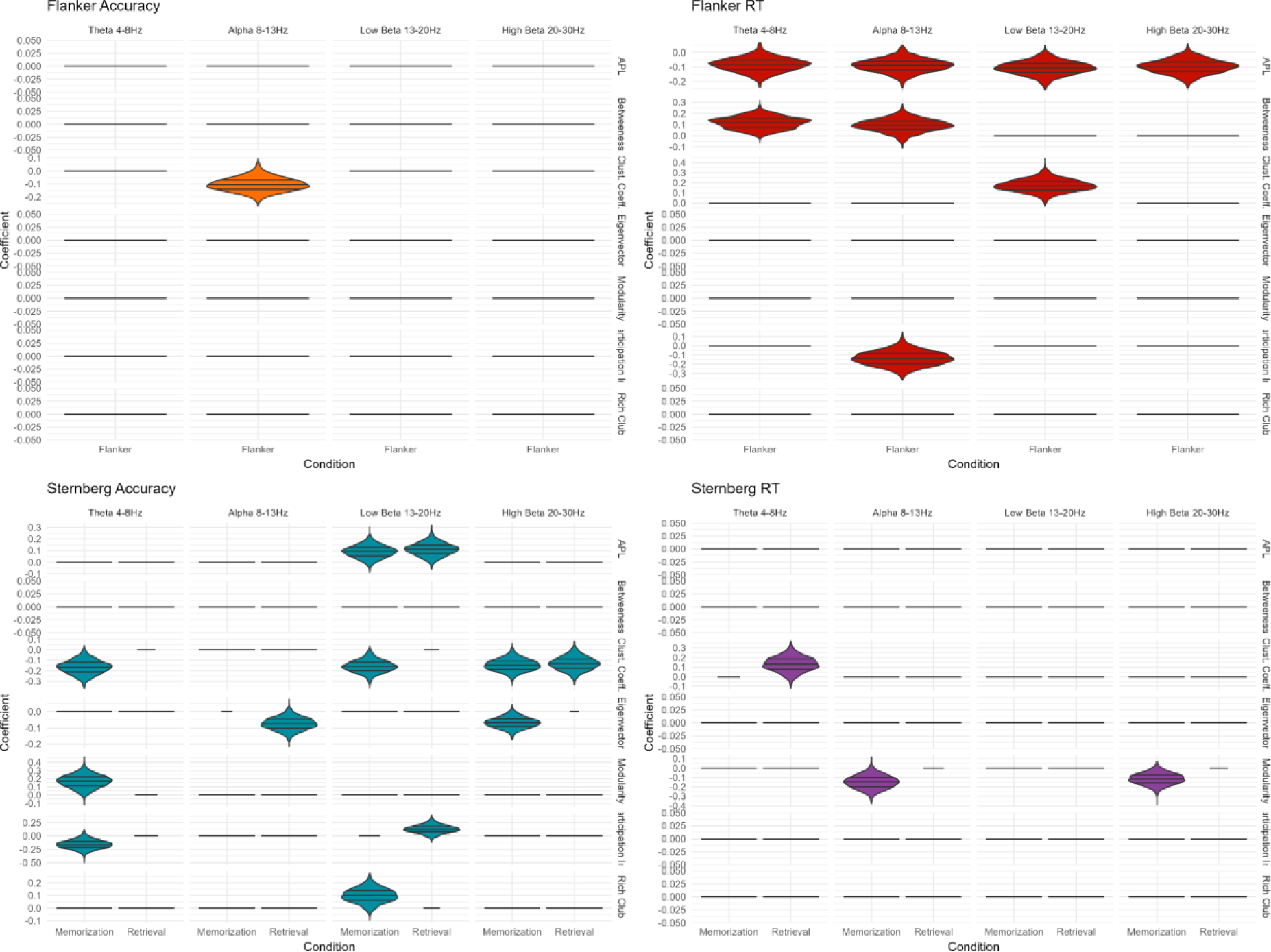
The significant effects of reconfiguration between resting state and cognitive state on behavioral measures at threshold 0.5. The x-axis represents the EEG condition, while the y-axis displays the regression coefficients of network measures on behavioral performance. The frequencies are visualized column wise, and the network measures are arranged rowwise. Each plot corresponds to a specific behavioral measure, with the violin plot indicating the distribution’s 0.25, 0.5, and 0.75 quantiles. Horizontal lines indicate absence of significant effect and were set to 0. Horizontal lines indicate absence of significant effect and are set to 0.

**Table # 8.**
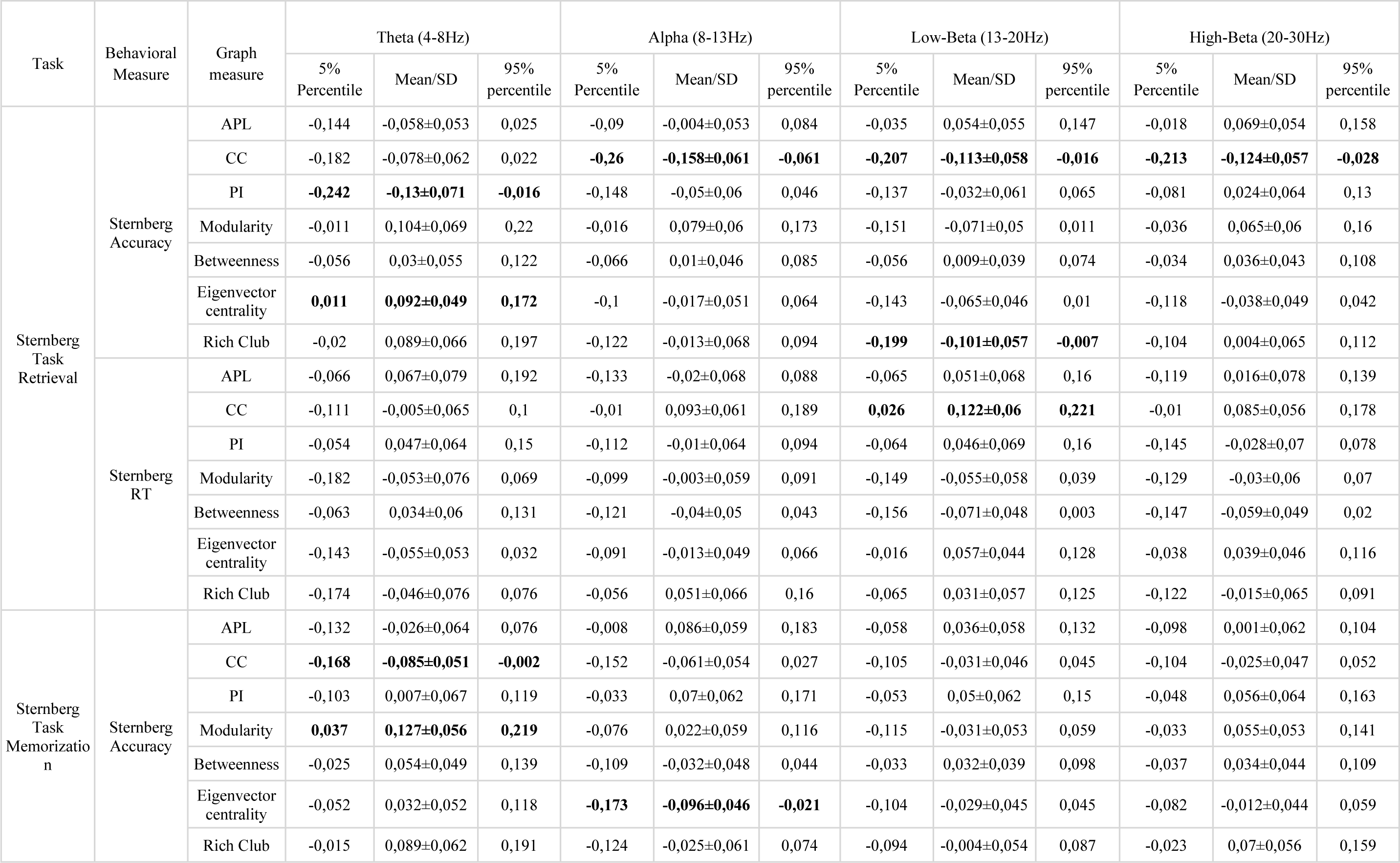
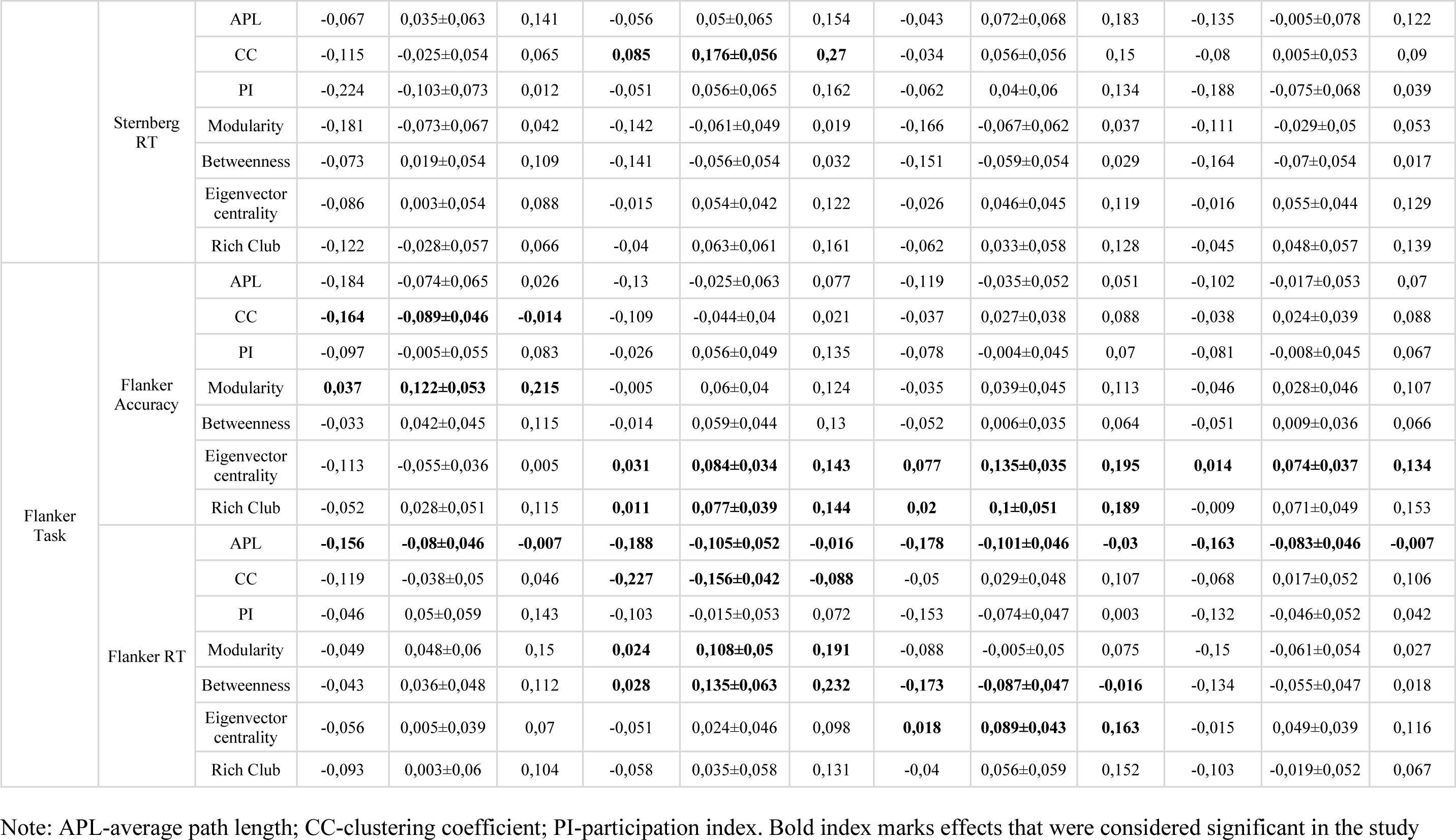
Regression coefficients for reconfiguration between resting state and cognitive tasks.

For the Sternberg task, neither path length nor betweenness centrality are related to test accuracy. The clustering coefficient in the theta rhythm is negatively associated with accuracy during the memorization phase. In the alpha band and both beta bands, it is similarly associated with the retrieval phase. Clustering coefficient is a sole predictor of response time with positive associations in alpha band and lower beta band. Eigenvector centrality positively affects memory at theta band but has a negative effect at alpha band: a negative impact on memory and a positive impact on response time. Accuracy is related to measures in theta band: positively with modularity during memorization phase and negatively with participation index during memory retrieval. In low beta band lower values of rich club coefficient during retrieval are associated with higher task accuracy.

For the Flanker task, path length is negatively correlated with response time across all frequency ranges (theta, alpha, low and high beta), indicating that greater reconfiguration towards enhancing information exchange between the resting state and the Flanker task is associated with longer reaction times. Centrality has varying effects on response time at different frequency ranges. In alpha band centrality positively affects response time, while in the low beta band the effect is negative. Clustering negatively correlates with correctness in theta band and with response time in alpha band, meaning that a higher level of clustering is associated with lower correctness and longer response times. Eigenvector centrality positively influences correctness in alpha, low and high beta bands, suggesting that a higher level of eigenvector centrality compared to resting state at these frequencies is linked to higher task correctness. Modularity has a positive impact on task correctness in theta band and on response time in alpha band, meaning that a higher level of modularity compared to resting state is associated with higher task correctness. Participation coefficient does not directly relate to test results in the Flanker task. The rich club is associated with task correctness in alpha and low beta bands in the Flanker task, with the higher level of rich clubs linked to higher task correctness at these frequencies.

The reconfiguration processes of brain networks seem to exhibit unique characteristics for each behavioral measure, but there are shared features across the Sternberg and Flanker tasks. The impact of the clustering coefficient on accuracy is a commonality; in the Flanker task, it specifically affects the theta band, whereas for the Sternberg task, it’s not restricted to a particular frequency band. Modularity in theta range also influences accuracy in both tasks, suggesting that network segregation into subnetworks is a general asset for cognitive performance. In the Sternberg task, increased clustering in the alpha and lower beta bands also correlates with longer response times, which is also true for the Flanker task.

For the Flanker task, however, accuracy is mostly influenced by eigenvector centrality across a range of frequencies, as well as by the rich club coefficient in alpha and low beta bands. This suggests that the roles of influential nodes and tightly interconnected hubs are critical for accurate attentional processing and inhibitory control. On the other hand, Sternberg task accuracy is primarily linked to the clustering coefficient, which seems to impact the frequency spectrum more broadly, indicating that localized network connectivity plays a significant role in working memory performance. Response time in the Flanker task is associated with network path lengths and measures of centrality and modularity, particularly in the alpha and low beta bands.

### Effect of functional connectivity at lower thresholds

Since the choice of a threshold value for removing spurious graph connections is arbitrary and there are no established guidelines for choosing this value, it is prudent to conduct an analysis for several thresholds. The threshold of 0.5 was chosen because our previous research (Zakharov et al., 2020) demonstrated that connectivity values obtained at this threshold are associated with cognitive characteristics. The comparison of thresholds shows that including weaker connections in the network indeed leads to additional associations with cognitive characteristics. The results reported in this section are presented in Figures 4, 5, 6 and Appendix #1.

**Fig 4.**
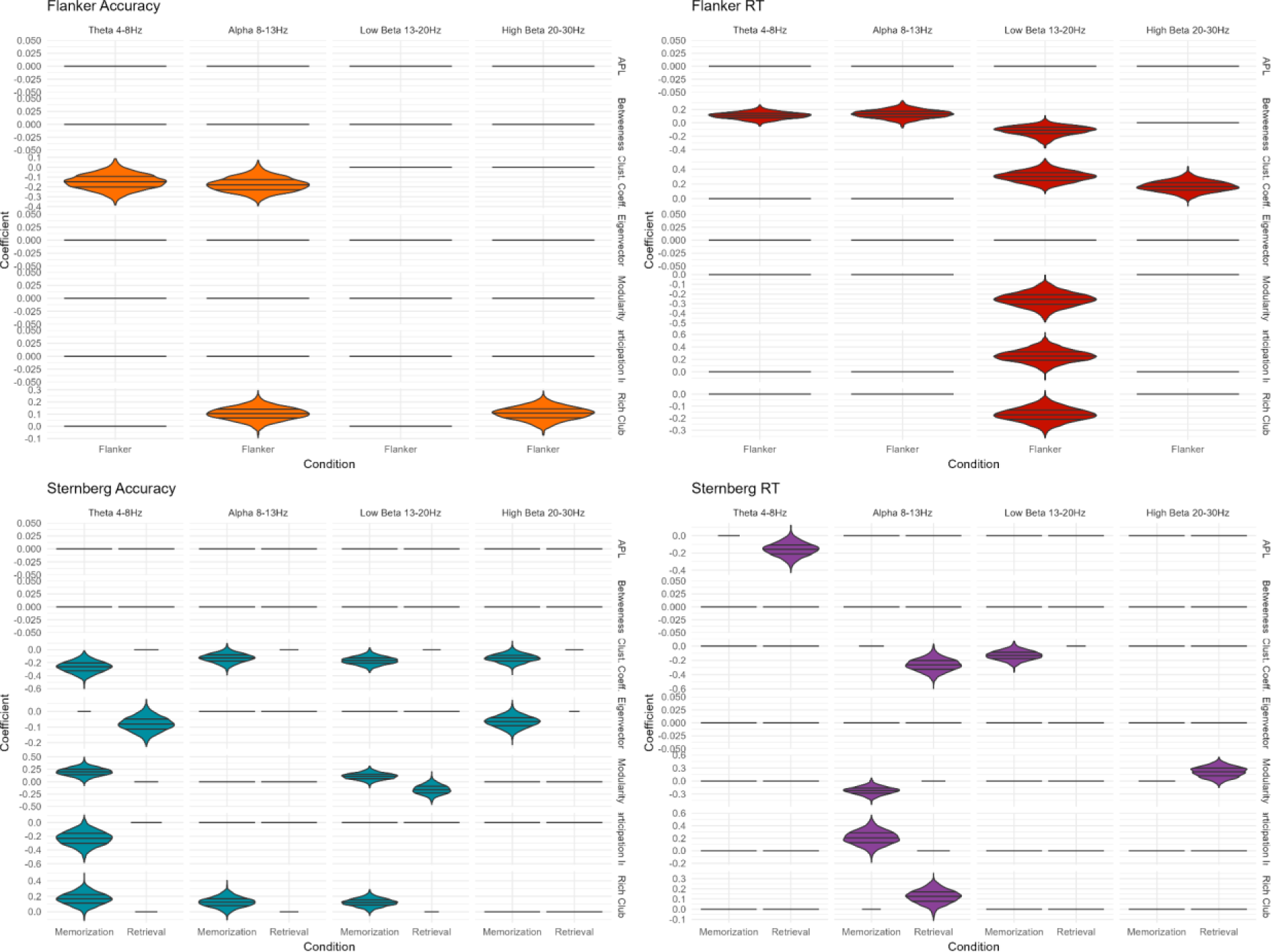
The significant effects of network measures during cognitive tasks on behavioral measures at threshold 0.5. The x-axis represents the EEG condition, while the y-axis displays the regression coefficients of network measures on behavioral performance. The frequencies are visualized column wise, and the network measures are arranged rowwise. Each plot corresponds to a specific behavioral measure, with the violin plot indicating the distribution’s 0.25, 0.5, and 0.75 quantiles. Horizontal lines indicates absence of significant effect and are set to 0.

**Fig 5.**
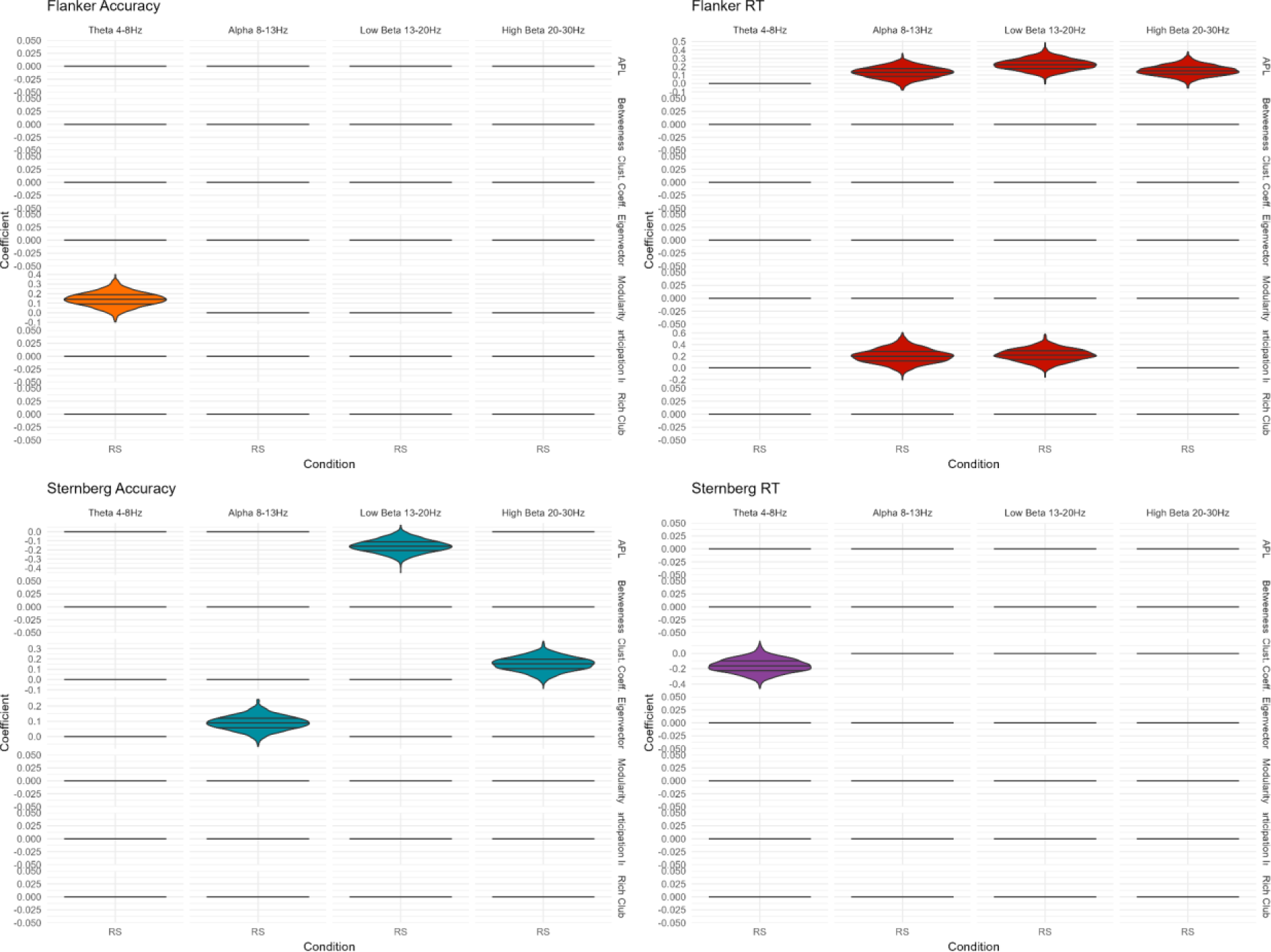
The significant effects of network measures during resting state on behavioral measures at threshold 0.5. The x-axis represents the EEG condition, while the y-axis displays the regression coefficients of network measures on behavioral performance. The frequencies are visualized column wise, and the network measures are arranged rowwise. Each plot corresponds to a specific behavioral measure, with the violin plot indicating the distribution’s 0.25, 0.5, and 0.75 quantiles. Horizontal lines indicates absence of significant effect and are set to 0.

This effect is most evident for accuracy in the Sternberg task and reaction time in the Flanker task during task execution. The Flanker task accuracy shows individual additional associations. In the Sternberg task, enhanced accuracy is linked to a reduced clustering coefficient across frequencies during the memorization phase, indicating less local interconnectivity. Additionally, increased modularity and rich club coefficients in the theta band, along with a decreased participation index in the same band, are also associated with higher accuracy. In the low beta band, accuracy correlates with heightened modularity and rich club connectivity during memorization. As for response times, they are tied to network modularity and participation index in the alpha band during memorization, as well as to average path length within the theta band and modularity in the high beta band. In the Flanker task, higher accuracy is associated with a lower clustering coefficient in the theta and alpha bands and with rich club connectivity in the alpha and high beta bands. When considering response time, new associations emerge with node centrality (betweenness centrality) in the theta, alpha, and low beta bands, and with nearly all measured network metrics in the low beta band.

In the resting state, an increase in Flanker task accuracy correlates exclusively with heightened modularity within the theta frequency band. Conversely, response time is affected by an increase in average path length in the alpha and both beta frequency ranges, as well as higher participation coefficients in the alpha and lower beta ranges. The determinants for accuracy in the Sternberg task are more dispersed: improved accuracy is linked to a shorter average path length in the lower beta band, an elevated clustering coefficient in the high beta band, and greater eigenvector centrality in the alpha band. As for response time in the Sternberg task, it is associated only with a reduced clustering coefficient within the theta frequency range.

The contrast between the resting state and cognitive task performance reveals a more extensive array of associations with certain cognitive measures. Specifically, accuracy in the Flanker task and response time in the Sternberg task are each associated with a limited number of network characteristics: for the Sternberg task, response time correlates with clustering coefficient in the theta band and modularity in the alpha and high beta bands. In contrast, accuracy in the Sternberg task and response time in the Flanker task exhibit a broader range of associations with network reconfiguration and behavioral outcomes. Sternberg task accuracy is linked to a host of network metrics, including a shorter average path length in the low beta range, clustering coefficients across theta and both beta bands, eigenvector centrality in both the alpha and high beta bands, modularity and participation coefficients in the theta band, as well as participation and rich club coefficients in the low beta range. Flanker task response time, meanwhile, is associated with a reduction in average path length across all frequency bands, an uptick in betweenness centrality in the theta and alpha bands, a rise in clustering coefficient in the low beta band, and a decreased participation index in the alpha band.

The results indicate that weak connections in the neural network play a significant role in cognitive functions, as demonstrated by their associations with performance on cognitive tasks. The inclusion of weaker connections in the analysis has revealed additional and meaningful associations with cognitive characteristics that might be overlooked when only stronger connections are considered. This highlights the importance of the brain’s full network, including its more subtle and less prominent features, in supporting cognitive processes. In the context of the Sternberg and Flanker tasks, which measure different cognitive abilities, the weak connections seem to contribute to the nuances of cognitive performance. For example, the finding that path lengths and the reconfiguration between background and test states are associated with the accuracy of the Sternberg task suggests that the integration of information across different regions of the brain, facilitated by these weak connections, is crucial for successful memory encoding and retrieval (Cohen & D’Esposito, 2016; Wang et al., 2021). Similarly, the positive association of the eigenvector centrality with reaction time in the Flanker task highlights those nodes which are weakly connected to many other nodes can influence cognitive processing speed. These weakly connected nodes might serve as crucial points of filtering the incoming information, enabling noise-free communication across the network. The negative associations observed between the clustering coefficient during memorization and weak connections across all frequency ranges indicate that a more segregated network might impede the ability to store information in the brain networks. This could be due to an over-reliance on local processing at the expense of broader network integration. Furthermore, the modularity’s association with the memorization phase suggests that a balance between segregated and integrated processing is crucial for memory formation. High modularity indicates a well-defined community structure within the network, which is beneficial for certain cognitive tasks, but too much segregation could hinder the kind of network-wide coordination required for others.

## 4. Discussion

In our work, we focused on comparing the Flanker and Sternberg tasks to identify the topological features of networks associated with successful performance in each, as well as to investigate the role of network reconfiguration between rest and cognitive load states. We assume that both processes will have a common core of network-related processes responsible for the general cognitive functioning, and a set of features unique for each behavioral measure.

Our investigation has unearthed several neural mechanisms that persist across different cognitive scenarios. The clustering coefficient in the alpha frequency band emerges as a common factor influencing both accuracy and response times in both the Flanker and Sternberg tasks. Modularity within the theta band and the rich club coefficient in the alpha band are linked to three out of four cognitive measures, with the exception being the response time in the Flanker task. Similarly, the participation coefficient and clustering coefficient in the low beta band show relevance to three cognitive measures, excluding the response time in the Sternberg task.

Nevertheless, the majority of the connections between network measures and cognitive performance are largely distinct between the Sternberg and Flanker tasks, and are specific to either accuracy or response time. Within the Flanker task, the neural processes underpinning accuracy appear to operate independently of those affecting response time. The pattern of associations suggests a transition in the frequency bands related to task performance, with accuracy associations moving from the beta band to the theta and alpha bands. Conversely, response time associations predominantly present in the alpha band during rest seem to shift towards the low beta range during the task. In contrast, the Sternberg task exhibits associations between cognitive performance measures—both accuracy and response time—and a more diverse array of frequencies and network measures, largely during the memorization phase. The Flanker task’s connections are more nuanced and may be highly specific, particularly as response time for this task is nearly exclusively linked to a singular frequency band.

Our results support the notion of inhibition being related to the more segregated network (Cole et al., 2013; Fair et al., 2012; Rubia et al., 2014) and working memory related to the more distributed network (Avery et al., 2020; Murphy et al., 2020) []. The Flanker task, which assesses inhibition, shows a different, more segregated, pattern. Task performance appears to be influenced by the individual’s general brain network organization at rest, suggesting a reliance on a broader range of cognitive processes rather than a focused, task-specific network [Cole et al., 2014]. In contrast, the Sternberg task, which is heavily reliant on working memory, within-task ability to induce a broader network might be important for recruitment of stimuli-specific brain areas (Ren et al., 2019) or particular processes in the working memory (Ekman et al., 2016). Response time seems to be influenced by the network’s ability to quickly reconfigure and efficiently utilize its resources. Faster response times may be facilitated by a network that can rapidly adjust its connectivity patterns (Gbadeyan et al., 2022). This dynamic reconfiguration may allow the network to bypass less efficient routes and engage in more direct, high-speed communication channels when a rapid response is required. The efficiency of these mechanisms is likely reflected in the balance of network attributes, with certain frequency bands responsible for modulation of these processes.

Effects related to frequency bands were found for all the investigated ranges. Alpha and theta bands, which are often linked to attention and encoding processes, are related to both cognitive tests. Thus, results might be related to facilitation of efficient information retention and retrieval in working memory (Herweg et al., 2020; Pavlov & Kotchoubey, 2022; Riddle et al., 2020; Wianda & Ross, 2019), top-down control as required to resolve cognitive conflict and filter out irrelevant stimuli (Avital-Cohen & Tsal, 2016; Buschman & Miller, 2007; Jensen & Mazaheri, 2010), and integration of information across distributed brain regions during attention-demanding tasks (Cavanagh & Frank, 2014). Alpha oscillations are involved in attention by gating sensory inputs and suppressing non-essential information, thus facilitating focused cognitive activity (Jensen & Mazaheri, 2010). The association of both beta band’s with the Sternberg task execution suggests that higher frequency neural oscillations might be involved in the active maintenance and manipulation of information in working memory (Engel & Fries, 2010; Schmidt et al., 2019).

Our data suggest that weak connections play a key role in the cognitive processes underlying performance on the Flanker and Sternberg tasks, which may rest upon several potential mechanisms. In the Flanker task weak connections might support the diffuse neural communication necessary for the broad integrative processes of attentional control. These weak links could facilitate the rapid reassignment of cognitive resources across the brain’s network, allowing for flexible adaptation to the task’s inhibitory demands (Bertolero et al., 2018). For the Sternberg task, weak connections may underlie the task’s requirement for maintaining and manipulating information over short periods. These connections could be crucial for establishing transient communication channels between brain regions that are not typically strongly connected, but must coordinate during the memorization and retrieval phases (Santarnecchi et al., 2014). This might reflect the brain’s strategy to engage complementary pathways to enhance memory performance, ensuring that information is not bottlenecked within a few strong connections but is instead diffusely supported across the network (Bassett et al., 2011). Furthermore, our findings align with the theory that weak connections contribute to the stability and flexibility of neural networks (Pajevic & Plenz, 2012). By providing additional routes for neural signaling, weak connections may endow the network with enhanced resilience against localized disruptions, ensuring sustained cognitive function even in the face of potential perturbations (X. Liu et al., 2022; Nassi & Callaway, 2009). This added redundancy could be especially beneficial during high-demand cognitive tasks where the robustness of information transfer is paramount.

The Network Neuroscience Theory (Barbey, 2018) suggests that general intelligence (g) arises from the unique topology and dynamics of the human connectome, highlighting the role of brain network architecture in enabling flexible network reconfiguration during goal-directed behavior. It also posits that both strong and weak neural connections are crucial for intelligent information processing. Additionally, the theory distinguishes between the neural underpinnings of crystallized and fluid intelligence: crystallized intelligence relies on stable and readily accessible network states, while fluid intelligence is associated with network states that are more adaptable but less easily accessed.

Our extension of this framework to additional cognitive measures generally aligns with the original concept. Both working memory and inhibition depends on different attributes of network dynamic, indicating the brain’s flexible adaptation to the unique demands of various cognitive tasks.

The Flanker task relies on more segregated networks with stable, and possibly stronger, connections, which constitute the basis for an efficient system of filtering information. Conversely, the Sternberg task is based on distributed networks that involve the engagement of weaker connections. These weaker connections may function as a dynamic and malleable network, enabling the temporary maintenance and manipulation of information. They could provide additional pathways for information flow, thus preventing over-reliance on a limited set of stronger connections and ensuring a more distributed and resilient strategy for memory-related processes (Honey et al., 2007).

A small yet consistent core set of processes shared by both cognitive abilities has been identified, likely representing a foundational network active across various cognitive functions. This may constitute the brain’s default operational mode and provide a foundation for general cognitive capabilities. This finding resonates with the notion of a common infrastructure underpinning cognitive processing (Krienen et al., 2014; Mill et al., 2017; Sadaghiani & Wirsich, 2020).

Our results also indicate that working memory is contingent upon the same processes that underlie fluid intelligence, as portrayed in the network neuroscience theory of intelligence (Clark et al., 2017; Shipstead et al., 2016; Unsworth et al., 2014). This suggests that working memory and fluid intelligence may share common neural mechanisms, aligning with the process overlap theory (Kovacs & Conway, 2016). This theory proposes that the general factor of cognitive abilities, or ‘g’, is reliant on the overlap between cognitive processes. The presence of a stable core of processes, along with a flexible periphery that is unique to each cognitive function, might indicate that an individual’s cognitive abilities hinge on the reconfiguration of these peripheral processes around the central network dynamics. This conceptual framework underscores the interdependence of core and peripheral processes in determining cognitive performance, both of which are crucial for understanding the sources of individual differences in cognitive abilities.

Our study has several limitations. First, the number of intervals chosen to analyze brain functional connectivity was arbitrary, indicating that more precise research is needed to understand how cognitive performance is related to functional connectivity in the context of specific stimuli. Additionally, examining individual trajectories of brain dynamics could be beneficial, as the dynamics during the task were not evaluated in our study. Another limitation is the lack of validation of our findings across independent datasets. While our research focused on working memory and cognitive control, these areas are just a segment of the wide spectrum of cognitive processes. Our methodology involved analyzing network construction at two threshold levels; however, the interplay between strong and weak neural connections is complex and warrants further investigation. Although we applied a whole-brain connectivity approach, cognitive performance may also be closely related to the dynamics within specific localized intrinsic brain networks. Lastly, given the interrelation between working memory and cognitive control, future studies should consider investigating the cross-task relationships between functional connectivity and cognitive performance, as this could uncover intricate interactions that affect cognitive functions.

Overall, we investigated the neural correlates of working memory and cognitive control by analyzing the Flanker and Sternberg tasks. Our findings indicate that while these tasks share some common neural mechanisms, they largely exhibit distinct network characteristics that underpin cognitive performance. The Flanker task is linked to more segregated networks, in contrast to the Sternberg task, which depends on more integrated networks, underscoring the importance of network reconfiguration in cognitive processes. Furthermore, weak connections were found to be essential for cognitive flexibility and resilience. Our results align with the Network Neuroscience Theory of Intelligence, suggesting unique network properties associated with specific cognitive functions. However, we propose that a combination of common and distinct processes is key to understanding individual differences in cognitive abilities.

