## Appendix 1 for "Task-specific topology of brain networks supporting working memory and inhibition"

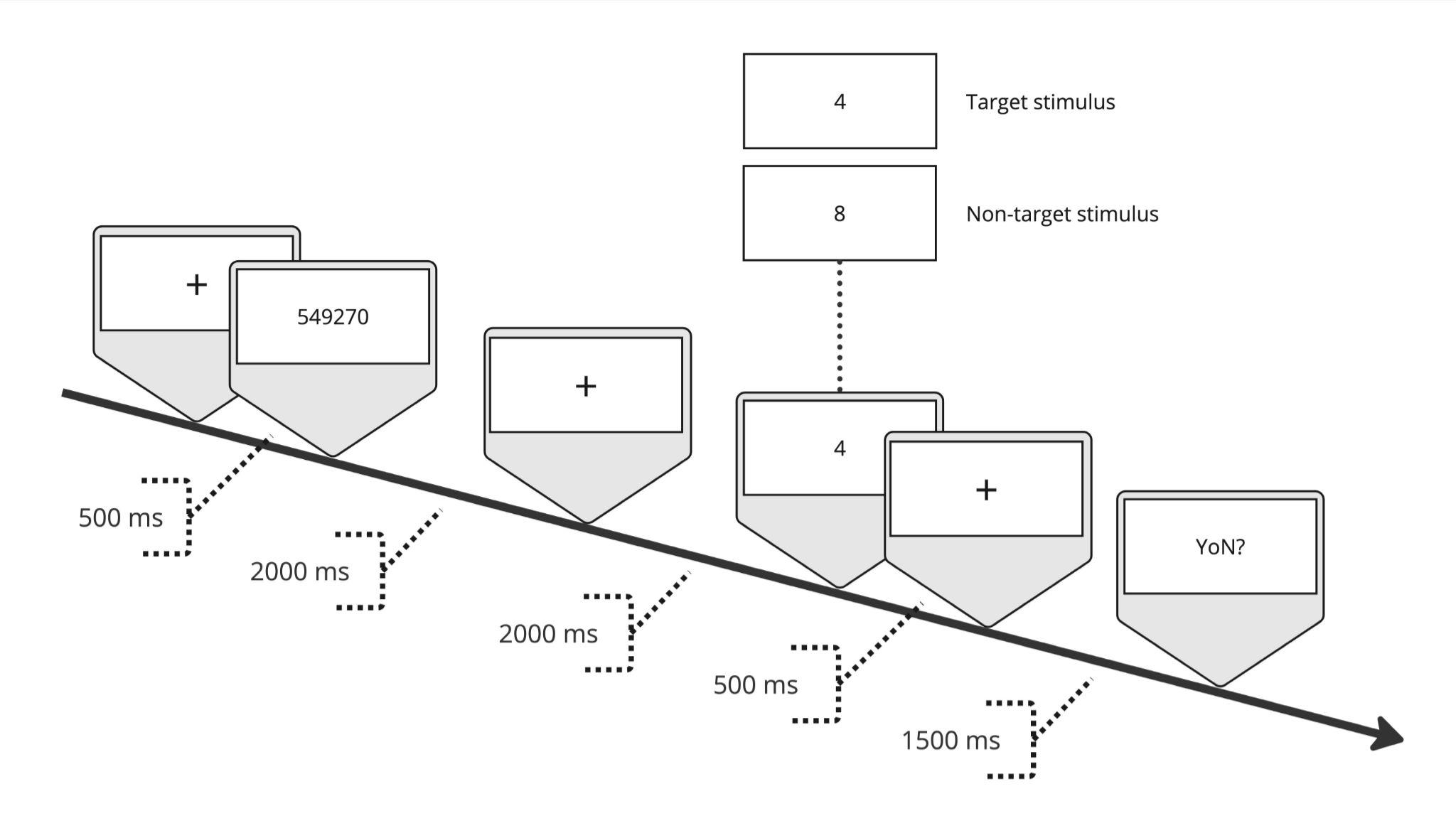

Fig1. Structure of the Sternberg task used in the study

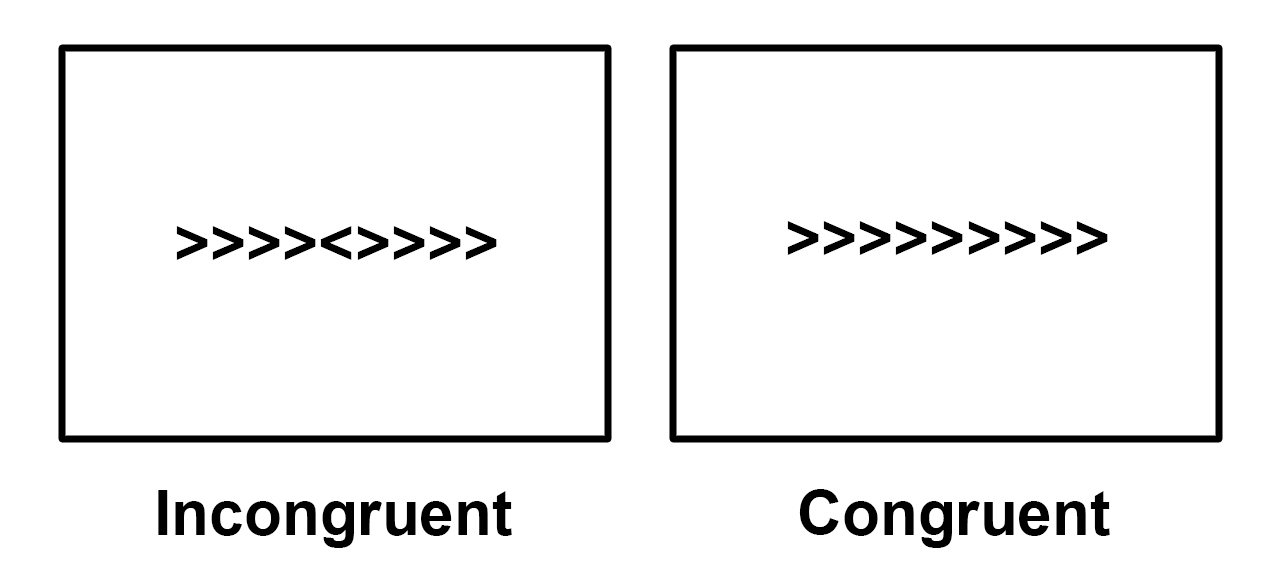

Fig2. Structure of the Flanker task used in the study. Source:

https://andysbrainbook.readthedocs.io/en/latest/fMRI_Short_Course/fMRI_02_ExperimentalDesign.html
